# Human astrocytes and microglia show augmented ingestion of synapses in Alzheimer’s disease via MFG-E8

**DOI:** 10.1101/795930

**Authors:** Makis Tzioras, Michael J.D. Daniels, Caitlin Davies, Paul Baxter, Declan King, Sean McKay, Balazs Varga, Karla Popovic, Madison Hernandez, Anna J. Stevenson, Jack Barrington, Elizabeth Drinkwater, Julia Borella, Rebecca K. Holloway, Jane Tulloch, Clare Latta, Jothy Kandasamy, Drahoslav Sokol, Colin Smith, Veronique E. Miron, Ragnhildur Thora Karadottir, Giles E. Hardingham, Christopher M. Henstridge, Paul M. Brennan, Barry W. McColl, Tara L. Spires-Jones

## Abstract

Synapse loss correlates with cognitive decline in Alzheimer’s disease (AD). Data from mouse models suggests microglia are important for synapse degeneration, but direct human evidence for any glial involvement in synapse removal in human AD remains to be established. Here we observe astrocytes and microglia from human brains contain greater amounts of synaptic protein in AD compared to non-disease controls, and that proximity to amyloid-β plaques and the *APOE4* risk gene exacerbate this effect. In culture, mouse and human astrocytes and primary mouse and human microglia phagocytose AD patient-derived synapses more than synapses from controls. Inhibiting MFG-E8 function rescued the elevated engulfment of AD synapses by astrocytes and microglia without affecting control synapse uptake. Thus, AD promotes increased synapse ingestion by human glial cells via an MFG-E8 opsonophagocytic mechanism with potential for targeted therapeutic manipulation.

**One-Sentence Summary:** Glial cells ingest synapses in Alzheimer’s disease and antibody treatment reduces this ingestion in cultured human cells.

## Introduction

Alzheimer’s disease (AD) is a neurodegenerative disease characterised by progressive cognitive decline and the accumulation of two pathological protein aggregates in the brain, amyloid-β (Aβ) plaques and phosphorylated tau tangles ^1,2^. Oligomeric forms of Aβ and tau are toxic to synapses and contribute to synapse loss ^3^. In turn, synapse loss is the strongest pathological correlate of cognitive decline in AD ^4,5^, however no therapies have been developed to effectively halt the degeneration and loss of synapses in humans. Controlled synapse elimination is an important aspect of the developing brain and this process is aided by glial cells in the brain, specifically astrocytes and microglia ^6–10^. This controlled synaptic refinement, or pruning, relies on multiple pathways, including the complement system ^7^, IL-33 ^11^ and epigenetic factors ^12^. Interestingly, it is hypothesised based on data from animal models that glial cells re-activate pathways involved in developmental synaptic pruning in a pathological manner during AD. For instance, in animal models of AD, microglia and astrocytes engulf more synapses both in response to Aβ pathology ^13–16^ and tau ^17,18^. The upregulated synaptic engulfment is often accompanied by cognitive deficits in mice, and importantly, when synaptic engulfment is blocked, there is a rescue of this cognitive deficit ^13,14,19^. This suggests that not all synapses that are removed by microglia are degenerating, and highlights an important therapeutic window for rescuing healthy synapses from being lost. At the moment, we still don’t know whether any of these findings translate to human brains. There is a paucity of human studies, which may in part contribute to the lack of effective drugs in trial or approved for AD^20^.

## Results

To date, there have been no quantitative studies measuring synaptic ingestion by glia in human brain comparing between AD and non-disease control cases. Here, we examined human post-mortem tissue from individuals with late-stage AD (n=31), age-matched control cases (n=19), and midlife control cases (n=9) without neurological disorders in two brain areas, the inferior temporal lobe (temporal cortex, BA20/21) which contains substantial Aβ and tau pathology, and the primary visual cortex (occipital cortex, BA17) which is affected later in dementia and contains less pathology than temporal cortex even at end stages of disease (information for human cases found at Table S1).

To determine whether astrocytes ingest synapses in human brain, we examined co-localisation between GFAP and Syn1 (Figure 1A) and observe a 2.1fold increase in AD compared to age-matched control brain (Fig. 1B) indicating astrocytes ingest more synapses in AD than control (p=0.0032). Further, there is a 2.7-fold increase in the volume occupied by synapses and GFAP between mid-life and healthy ageing (p<0.0001), indicating that astrocyte ingestion of synapses increases during age and further increases in AD (p<0.0001) (Figure 1B). Astrocytes ingest more synapses in the proximity of Aβ plaques in BA20/21 (p=0.0004) but not in BA17 in AD brain (Figure 1C and Figure 1 S1), and also astrocytes in *APOE4* carriers, who are more at risk of developing AD and who undergo more synapse loss ^21,22^, ingest more synapses than astrocytes in *APOE3* individuals (p=0.0003) (Figure 1D and Figure S1). Astrocyte burden was increased in AD brain and was higher in *APOE4* carriers as expected (plots not shown, data included in supplemental data spreadsheet). Moreover, there is more colocalisation of astrocytes and synapses in BA17 than BA20/21 (p<0.0001) (Fig. 1E), which occurs both in aged control and in AD cases (Fig S1B). When ingestion was normalised to GFAP burden, there was still a significant effect of cohort (p<0.0001) with both AD and aged controls having more synapse colocalization with astrocytes than mid-life controls in post-hoc tests (Figure S1C). Interestingly, women had significantly more synaptic ingestion by astrocytes when normalized to GFAP burden (p=0.02).

**Figure 1.**
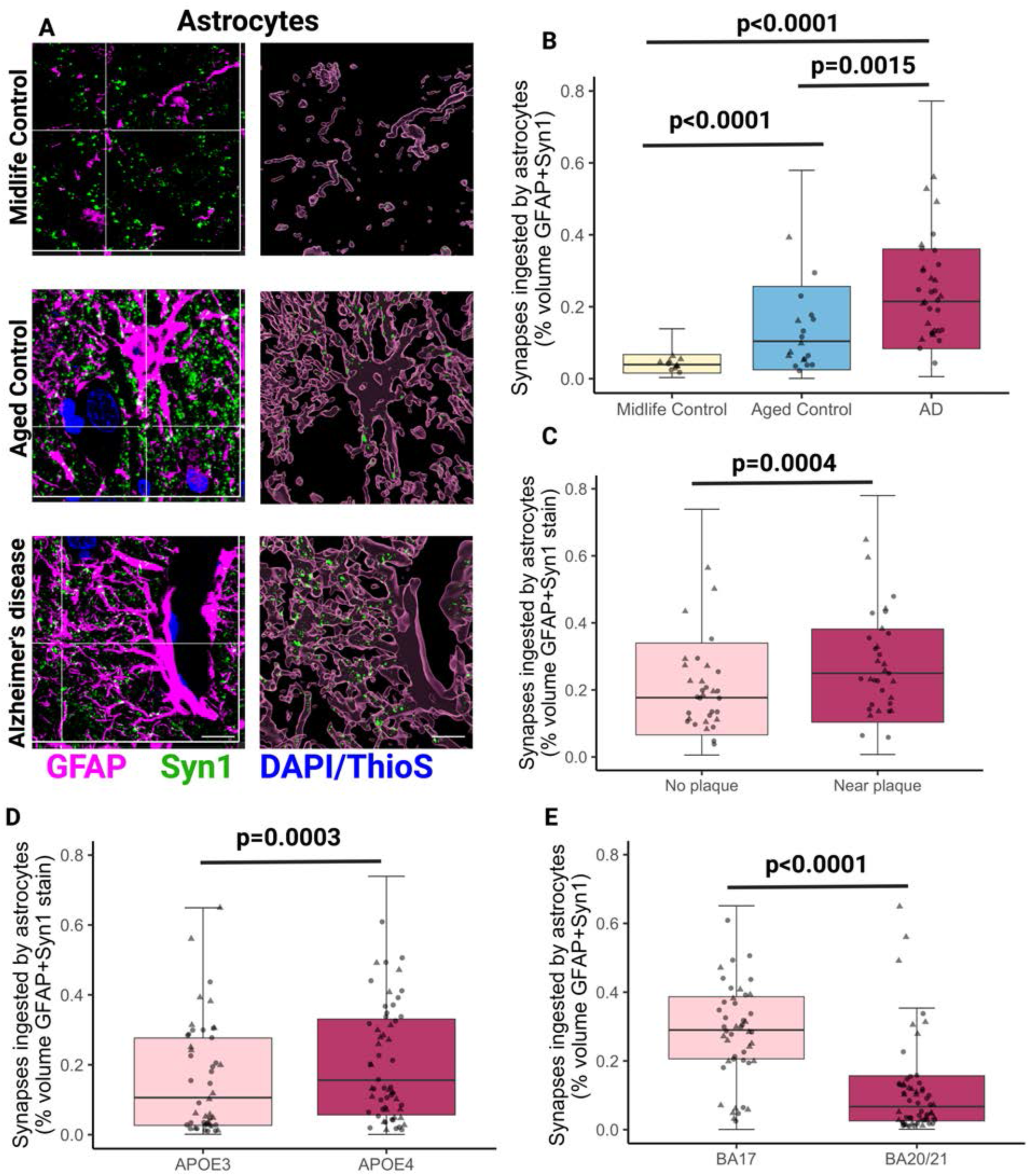
Astrocytes ingest more synapses in Alzheimer’s disease compared to midlife and aged controls. (A) Confocal images of immunostaining with orthogonal views (left) and three-dimensional reconstructions of stacks (right) of midlife control, aged control and Alzheimer’s disease showing ingestion of synapsin 1 (Syn1, green) by astrocytes (GFAP, magenta) in human brain sections. Aβ plaques and nuclei are counterstained with Thioflavin S and DAPI, respectively, shown in blue. Only ingested Syn1 is shown in 3D Imaris reconstructions. Representative images are from BA17. (B) Quantification of synapsin 1 ingested by GFAP-positive astrocytes showed significantly increased levels in AD compared to both midlife controls (post-hoc Tukey corrected tests after linear mixed effects model t=7.59, p<0.0001) and aged controls (t=3.38, p=0.0015), and also between midlife controls to aged controls (t=4.60, p=0.001). Data here are pooled from both BA17 and BA20/21. Males are represented by circles and females by triangles. (C) Statistically significant increase of synapsin 1 ingestion by astrocytes near Aβ plaques in AD cases (F[1,1198]=12.43, p=0.0004). (D) The APOE4 genotype was associated with an increase in synapsin 1 colosalisation inside GFAP-positive astrocytes (F[1,77.35]=14.31, p=0.0003). (E) BA17 (primary visual cortex) was associated with higher levels of synapsin 1 colosalisation inside GFAP-positive astrocytes compared to BA20/21 (inferior temporal cortex) (F[1,2226.80]= 1372.67, p=2.2×10^-16^). Statistical comparisons were made using ANOVA after linear mixed effects model on Tukey transformed data with case as a random effect and disease, brain region, APOE4 status, sex, and age as fixed effects. Untransformed data are presented in graphs B-E all data are included in box plots and case medians are shown in points. Scale bars represent 5μm.

We also assessed internalisation of synapses by microglia using Syn1 colocalisation with the microglial lysosomal marker CD68 (Figure 2A) which was also confirmed by super-resolution Airyscan microscopy (Figure S1D). CD68 labels both microglia and macrophages and in the cortical neuropil regions examined from our cases, we observe that CD68 positive cells were also positive for IBA1 and the microglial marker TMEM119 ^23^, Figure S1E). We found a significant increase in the volume of co-localisation between CD68 and Syn1 in AD compared to aged controls (p=0.0024) and in AD compared to mid-life controls (p=0.0096), suggesting increased synaptic ingestion by microglia in disease (Fig. 2B). The increased synaptic ingestion by microglia in AD was present in both BA17 and BA20/21 (Figure S1F). Unlike astrocyte ingestion, there was no increase in microglial ingestion of synapses between mid-life and healthy ageing (p=0.625) (Fig. 2B). Moreover, in AD brain samples, this ingestion was higher in the presence of Aβ plaques (p<0.0001) (Figure 2C), and the presence of the *APOE4* risk allele was also associated with increased synaptic ingestion by microglia in individuals with AD (p=0.02) (Figure 2D and Figure S1F). Similar to astrocytes, there is more colocalisation of microglia and synapses in BA17 than BA20/21 (Figure 1E and Figure S1F). As expected, we observe microgliosis in AD brain with a 1.9-fold increase in CD68 volume across both brain areas (plots not shown, data included in Supplemental Data spreadsheet 1). When synaptic colocalisation with CD68 was normalised to CD68 volume, no differences were seen between AD and control brain indicating that the increased colocalization (reflective of ingestion) is at least partly driven by microgliosis and/or microglial hypertrophy. These data show that microglia contain synaptic protein in human brain with more ingestion in AD, more near plaques, and more in APOE4 carriers.

**Figure 2.**
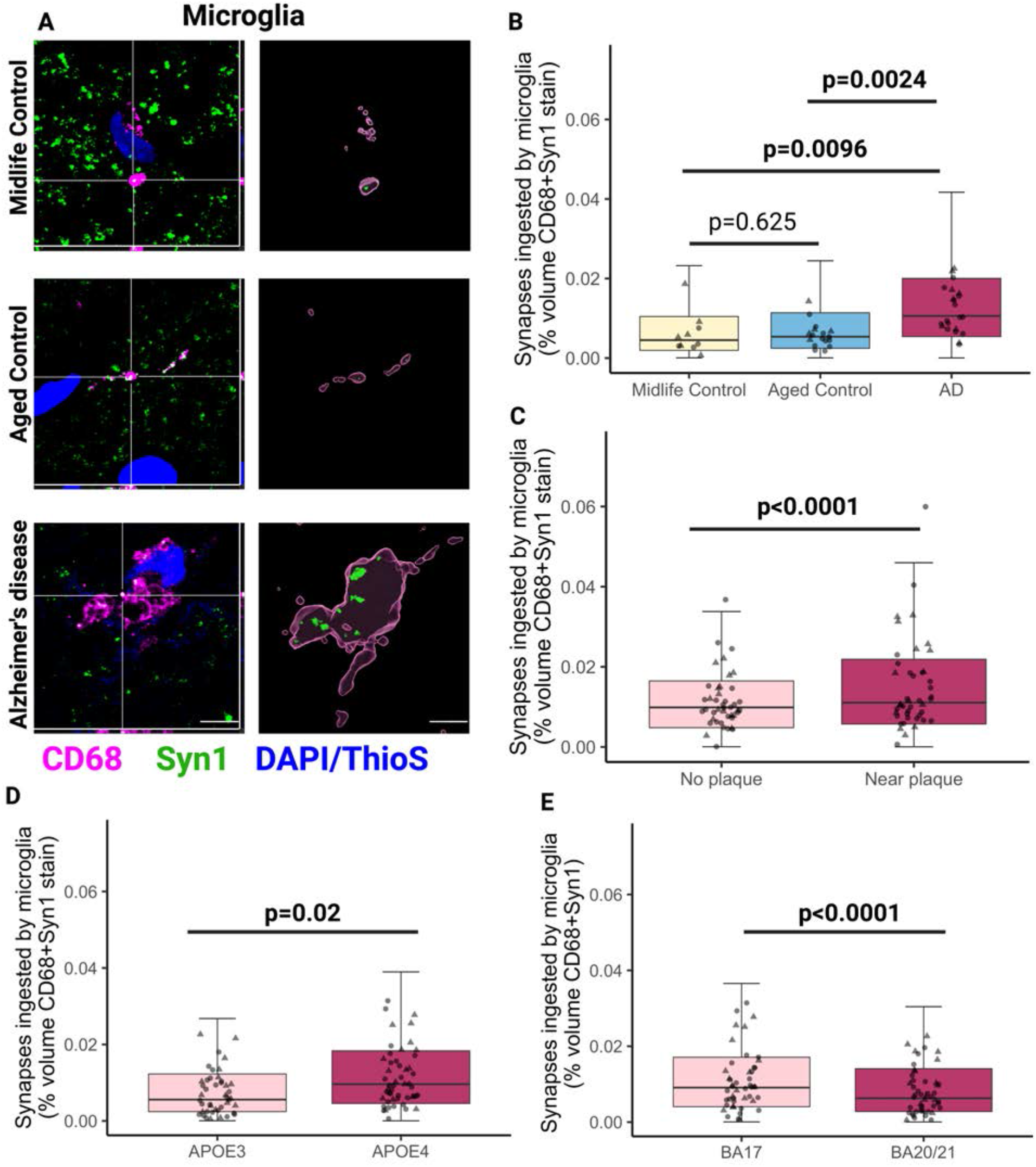
Microglia ingest more synapses in Alzheimer’s disease compared to midlife and aged controls. (A) Confocal images of immunostaining with orthogonal views (left) and three-dimensional reconstructions of stacks (right) of midlife control, aged control and Alzheimer’s disease showing ingestion of synapsin 1 (Syn1, green) by microglia (CD68, magenta) in human brain sections. Aβ plaques and nuclei are counterstained with Thioflavin S and DAPI, respectively, shown in blue. Only ingested Syn1 is shown in 3D Imaris reconstructions. Representative images are from BA17. (B) Quantification of synapsin 1 ingested by CD68-positive microglia showed significantly increased levels in AD compared to midlife (Tukey corrected post-hoc test t=3.56, p=0.0024) and aged controls (t=3.09, p=0.0096). No statistical difference was seen between midlife controls and aged control (t=0.93, p=0.62). Data here are pooled from both BA17 and BA20/21, sample size corrected by mixed effected linear model. Males are represented by circles and females by triangles. (C) Statistically significant increase of synapsin 1 ingestion by microglia near Aβ plaques in AD cases (ANOVA F[1,881.57]=20.74, p=6.01×10^-6^). (D) The APOE4 genotype was associated with an increase in synapsin 1 colosalisation inside CD68-positive microglia (F[1,42.86]=5.84, p=0.02). (E) BA17 (primary visual cortex) was associated with higher levels of synapsin 1 colosalisation inside CD68-positive microglia compared to BA20/21 (inferior temporal cortex) (F[1,1973.76]=67.82, p=3.42×10^-16^). Statistical comparisons were made using post-hoc Tukey test or ANOVA after linear mixed effects model on Tukey transformed data with case as a random effect and disease, brain region, APOE4 status, sex, and age as fixed effects. Untransformed data are presented in graphs B-E. Scale bars represent 5μm.

To further validate microglial and macrophage ingestion of synapses, we immunostained for the microglial-specific marker P2Y12 which confirmed synaptic colocalisation with this marker (Figure 3A). Since synapsin 1 can be observed in axons as well as synapses, we confirmed synaptic ingestion by both astrocytes and microglia using synaptophysin, a more specific presynaptic vesicle marker (Figure 3A). We also observe excitatory postsynaptic densities (PSD95) and inhibitory presynaptic proteins GAD65/67 within astrocytes and microglia (Figure S2A and B). The synaptic ingestion we observe in post-mortem samples could be due to specific phagocytosis of synapses from living neurons or more general clearance of dead or dying neurons by glia. To investigate this possibility, we co-stained synaptic protein synaptophysin, neuronal neurite protein MAP2, astrocytes, and microglia (Figure 3A). Similarly to our previous experiments, we observe significantly higher colocalisation of synaptophysin with both astrocytes (Figure 3B) and microglia (Figure 3C). Interestingly, no significant differences in P2Y12 burdens were observed between aged controls and AD individuals (p=0.1526, data not shown) suggesting the increased synaptic ingestion is not due to microgliosis. Triple colocalisation of MAP2 with synaptophysin and glia showed a 26-fold decrease in ingestion compared to synaptophysin alone (Figure 3D-E), suggesting synaptic ingestion occurs predominantly in the absence of neuronal death. We also did not observe axonal neurofilament ingestion by astrocytes and microglia (Figure S2C).

**Figure 3.**
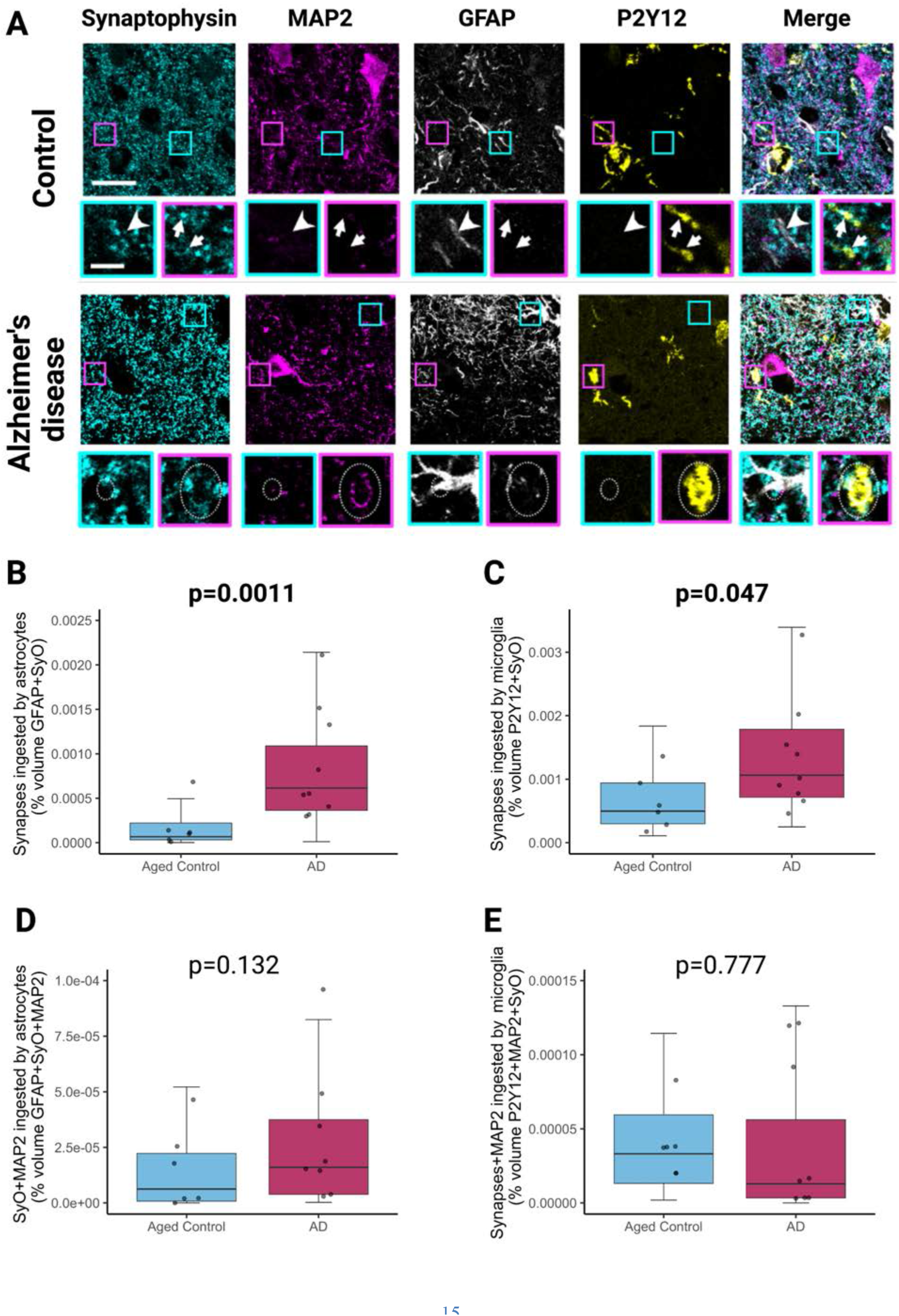
Astrocytes and microglia ingestion of synapses in the absence of neuronal neurite ingestion. (A) Staining presynaptic terminals with synaptophysin, neuronal neurites with MAP2, astrocytes with GFAP, and microglia with P2Y12 reveals synaptic ingestion by astrocytes (cyan boxes, arrowheads) and microglia (magenta boxes, arrows) in the absence of MAP2 staining in control and AD brain. In AD, MAP2 positive neuronal protein can also be observed in astrocytes (cyan boxes, dotted ovals) and microglia (magenta boxes, dotted ovals). Large panels are maximum intensity projections of confocal image stacks. Insets are single sections to demonstrate colocalization. Scale bars represent 20 μm in large panels, 5 μm in insets. (B) Quantification of synaptophysin (SyO) ingested by GFAP-positive astrocytes showed significantly increased levels in AD (n=9) compared to aged controls (n=6) (post-hoc Tukey corrected tests after linear mixed effects model F[1,13.03]=17.38, p=0.0011). (C) Quantification of synaptophysin (SyO) ingested by P2Y12-positive microglia showed significantly increased levels in AD (n=9) compared to aged controls (n=6) (post-hoc Tukey corrected tests after linear mixed effects model F[1,13.00]=4.8000, p=0.0473). (D) Quantification of synaptophysin (SyO) and MAP2 ingested by GFAP-positive astrocytes showed no significant differences between AD (n=9) and aged controls (n=6) (post-hoc Tukey corrected tests after linear mixed effects model F[1,13.01]=2.57, p=0.132). (E) Quantification of synaptophysin (SyO) and MAP2 ingested by P2Y12-positive microglia showed no significant differences between AD (n=9) and aged controls (n=6) (post-hoc Tukey corrected tests after linear mixed effects model F[1,12.98]=17.38, p=0.777).

Having shown in post-mortem human tissue that there is more synaptic protein located within microglia and astrocytes in AD compared to control brains, we next sought to determine whether synapses derived from human AD brains are more readily internalised by glia in dynamic *ex vivo* assays. We used snap-frozen samples from the temporal lobe (Brodmann area 38, or BA38) of control and AD cases to prepare synaptoneurosomes and synaptosomes in synapse-enriched fractions (SEFs), as previously validated by our lab^24^ and others^25^. Western blotting showed the synaptoneurosomes and synaptosomes had significantly higher levels of both synaptophysin (Sy38) and post-synaptic density 95 (PSD-95) in the SEFs compared to total brain homogenate, as well as a significant de-enrichment of the nuclear marker histone H3 (Figures S3A-C and S3E-F). Synaptic integrity was also validated by electron microscopy (Figure S3G). We observed lower levels of synaptophysin in AD brain homogenates and synaptoneurosomes compared to brains from age-matched non-demented controls, reflecting the known synapse loss that occurs in AD (Figure S3D).

Synaptoneurosomes and synaptosomes were conjugated to pHrodo-Red succinidyl ester, enabling visualisation of synapses internalised within the acidic phago-lysosomal compartment during phagocytosis ^26,27^. We challenged cultured GFAP-expressing human and mouse astrocytes (Figure S4A-B and Supplemental Videos 1 and 2) with pHrodo-Red labelled synaptoneurosomes from AD or control brain to confirm they phagocytose synapses (Figure 4A and B). We observe that cultured human astrocytes phagocytose synaptoneurosomes from AD brain both more (p=8.47×10^-4^) and faster (p=8.25×10^-14^) than those from controls (Figure 4C and D). Similarly, AD synaptosomes were also ingested more than control synaptosomes by human astrocytes (Fig S3H). Primary mouse astrocytes similarly ingested AD synapses both more (p=8.×10^-4^) and faster (p=2×10^-16^) than controls (Fig 4E-F). Importantly, phagocytosis of synaptoneurosomes was completely suppressed in astrocytes treated with cytochalasin D (CytD), a potent inhibitor of actin polymerisation which prevents phagocytosis (Figure 4C and E; Figure S4C-D).

**Figure 4:**
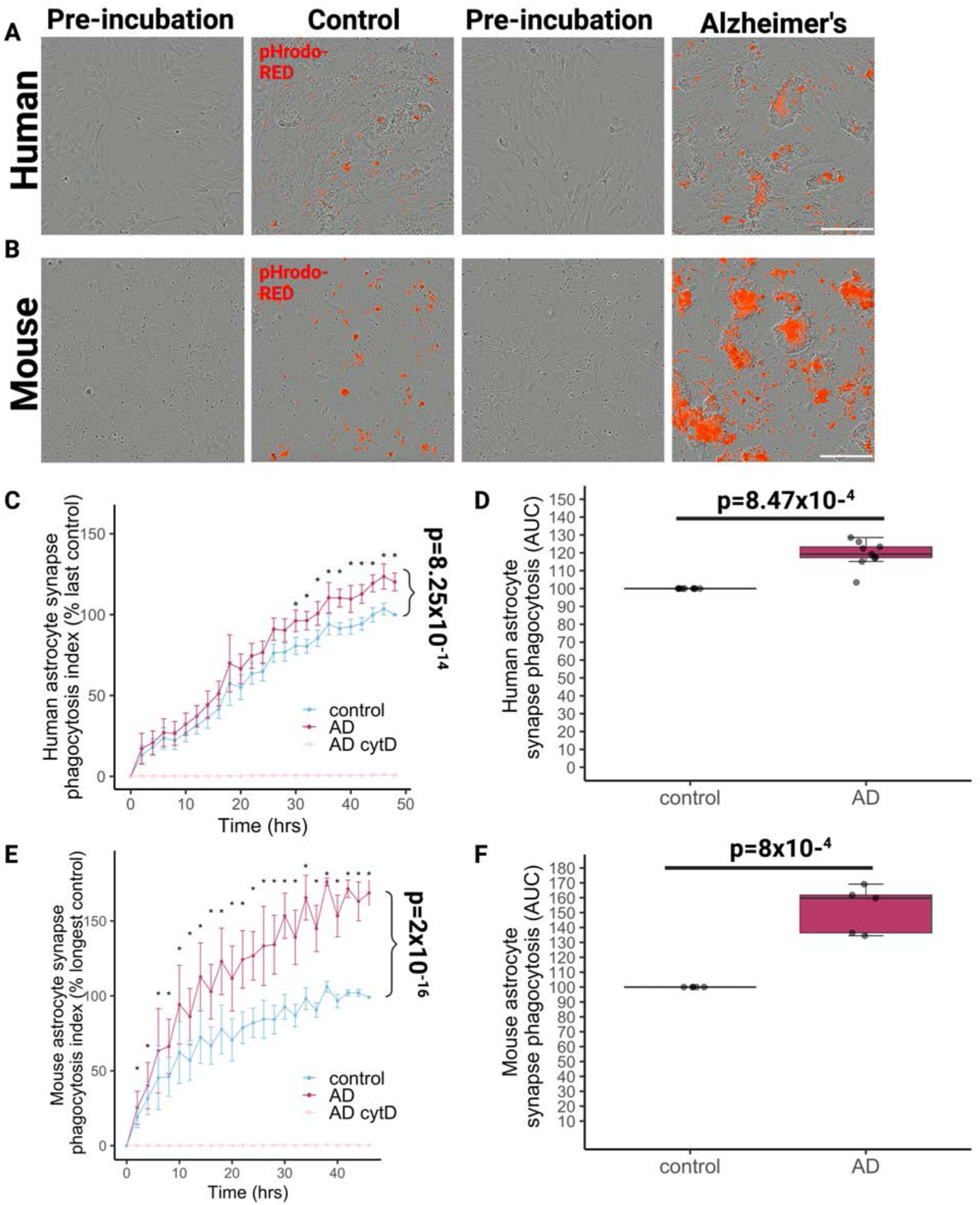
Human and mouse astrocytes ingest human AD synapses more than control synapses in culture. (A) Live-imaging assay of primary human foetal astrocytes shows astrocytes phagocytose pHrodo-Red labelled human synaptoneurosomes from control and Alzheimer’s disease (AD) brains after treatment (48 hours). Scale bar represents 200μm. (B) Live-imaging assay of primary mouse embryonic astrocytes shows astrocytes phagocytose pHrodo-Red labelled human synaptoneurosomes from control and Alzheimer’s disease (AD) brains after treatment (48 hours). Scale bar represents 200μm. (C) The phagocytosis index curves normalised to within experiment control at the last time point show that human astrocytes (n=8 independent replicates) phagocytose AD synapses more and faster than controls (ANOVA after linear mixed effects model with disease status of synapse donor and incubation time as fixed effects and experimental replicate as random effect, effect of disease status F[1,370.03]=50.25, p<0.0001, effect of time F[24,370.09]=85.45, p<0.0001). Asterisks represent significant post-hoc Tukey corrected tests between AD and control at the time points indicated. (D) Examining the area under the curve confirms that AD synapses are phagocytosed more by human astrocytes than control synapses (ANOVA after linear mixed effects model with disease status of synapse donor as fixed effect and experimental replicate as random effect F[1,9.60]=61.14, p=1.83×10^-5^). (E) Mouse astrocytes (E-F, n=5 independent replicates) also ingest AD synapses more and faster than controls (effect of disease of synapse donor F[1,163.06]=230.98, p<0.0001; effect of incubation time F[25,163.17]=28.40, p<0.0001; Asterisks represent significant post-hoc Tukey corrected tests between AD and control at the time points indicated. (F) Examining the area under the curve confirms that AD synapses are phagocytosed more by mouse astrocytes than control synapses (F[1,4.80]=54.65, p=0.0008). CytD which inhibits phagocytosis completely prevented synapse ingestion.

We next tested whether microglia also ingest AD-synapses differently than controls. The BV2 murine microglial cell line was used to validate phagocytosis of pHrodo-tagged synaptoneurosomes where clear uptake of synaptoneurosomes into acidic subcellular compartments was observed, while this was diminished by CytD (Figure S4E-F). We then tested uptake of synaptoneurosomes from control or AD brains by primary human microglia isolated from peritumoural tissue extracted during glioblastoma surgeries (n=11 patient donors). Microglia were isolated by immunomagnetic separation, as previously validated for mice by our group ^28^, and human cells were shown to express the microglia-specific marker TMEM119 (Figure 5A). Human microglia in culture were exposed to AD or control synaptoneurosomes labelled with pHrodo and imaged using live microscopy (Figure 5B-C Supplemental Videos 3 and 4). A significantly greater proportion of human microglia phagocytosed AD synaptoneurosomes compared to controls (p=0.027) and this process also occurred faster (p=2×10^-16^) (Figure 5D-E). There were no effects of sex, age, or resection brain region for the microglial surgical donors. We also observed similar results from adult primary microglia from non-diseased temporal cortex of a neurosurgical biopsy due to epilepsy (Figure S5A-C), as well as from human pluripotent stem-cell derived microglia-like cells (Figure S5D-H). Given the inter-person variability of human microglia, we also challenged adult primary mouse microglia with the human synaptoneurosomes. Similarly to the human microglia, murine microglia also ingested AD synaptoneurosomes both more (p=0.01) and faster (p<0.0001) than control synaptoneurosomes (Fig. 5F-H, effect of disease of synapse donor: F[1,2.88E21]=102.39, p<0.0001; effect of incubation time F[18,inf]=98.10, p<0.0001). Primary mouse microglia also preferentially ingested AD synaptosomes compared to control synaptosomes (Figure S3I). Interestingly, cultured astrocytes phagocytosed synapses more slowly than microglia with maximum phagocytosis occurring at around 60 minutes in microglia and around 48 hours in astrocytes. Lastly, we excluded the possibility that AD-derived synapses interfere with the phago-lysosomal compartment to account for this increase in ingestion by running degradation assays in cultured astrocytes and microglia showing effective degradation of both AD and control synaptoneurosomes (Figure S6A-C).

**Figure 5.**
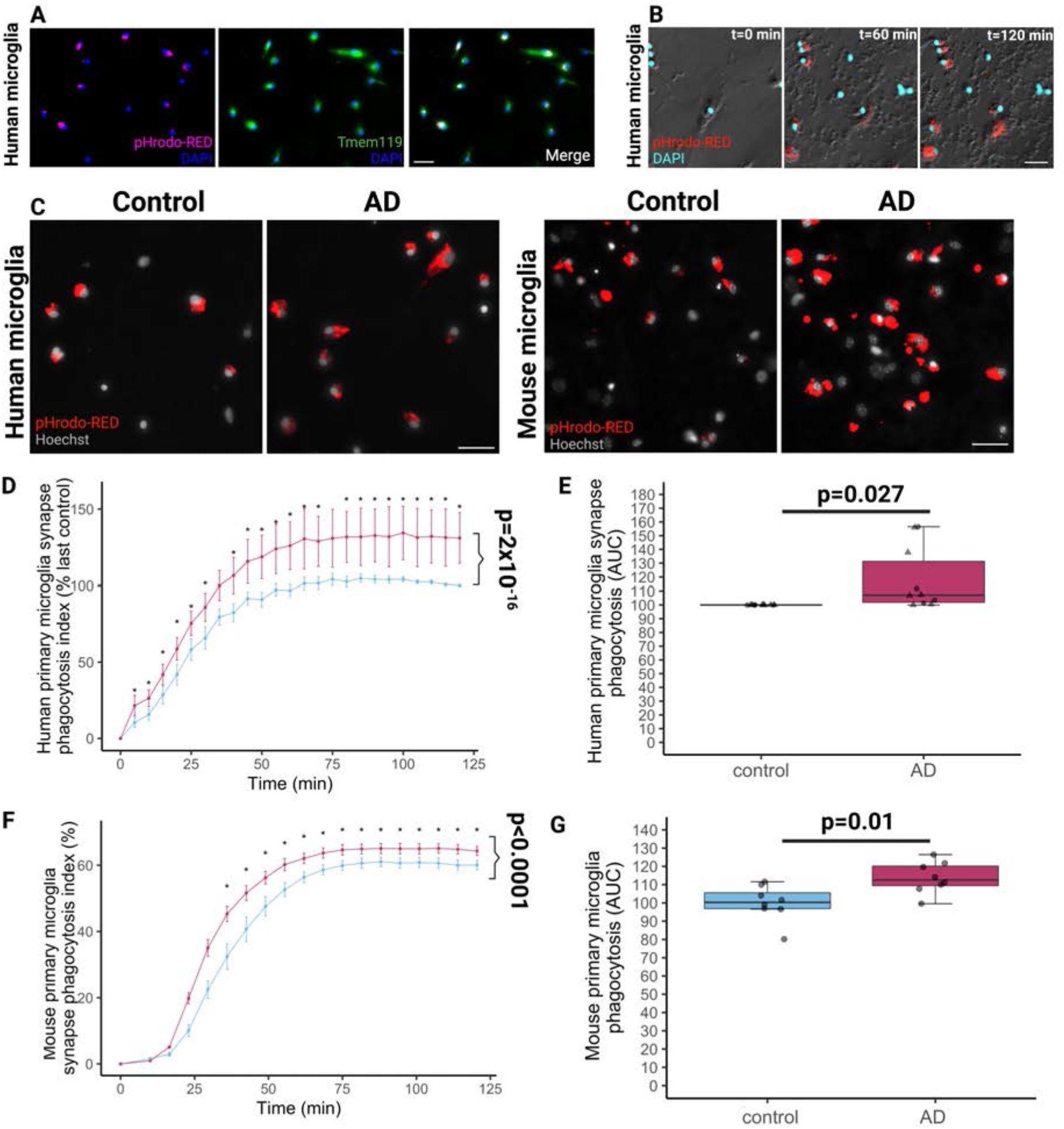
Human and mouse microglia ingest human AD synapses more than control synapses in culture. (A) Immunocytochemistry on fixed microglia grown from human peri-tumoural resected during neurosurgery shows they express the microglial specific marker TMEM119 and engulfed pHrodo-Red labelled human synaptoneurosomes when these were applied 2 hours before fixation. Scale bar represents 20μm. (B) Live imaging of human microglia (phase) with DAPI (cyan) and pHrodo-Red confirms live phagocytosis of human synapses over 2 hours. Scale bar represents 20μm. (C) Live imaging of human microglia with nuclear marker Hoechst and pHrodo-Red confirms that these primary cultured microglia engulf synapses derived from human AD and control brain. Scale bar represents 20μm. (D) Quantification of experiments from n=11 neurosurgical donors shows that the phagocytosis index normalised to the final value for the control condition for each experiment increases over time and that AD synapses are phagocytosed more and faster than control synapses (ANOVA after linear mixed effects model on square root transformed data with synaptoneurosome donor disease status, incubation time, neurosurgical brain region, microglial donor sex and microglial donor age as fixed effects and donor as a random effect, effect of disease status of synapse donor F[1,396.09]=122.74, p<2×10^-16^, effect of incubation time F[24,396.25]=119.49, p<2×10^-16^). (E) Quantifying the area under the curve of the phagocytosis index confirms more phagocytosis of AD than control synapses (* ANOVA after linear mixed effects model with synaptoneurosome donor disease status, incubation time, neurosurgical brain region, microglial donor sex and microglial donor age as fixed effects and donor as a random effect, effect of disease status of synapse donor F[1,12]=6.38, p=0.027). (F) Microglia grown from adult mouse brain phagocytose human synapses. Curves from n=8 mice show increased phagocytosis of AD compared to control synapses (ANOVA after linear mixed effects model with synaptoneurosome donor disease status and incubation time as fixed effects and mouse microglia donor as a random effect, effect of disease F[1,2.88E21]=102.39, p<0.0001, effect of incubation time F[18,inf]=98.10, p<0.0001). (G) Increased ingestion of AD synaptoneurosomes compared to control ones confirmed by an increased area under the curve in AD (* ANOVA effect of disease status of synapse donor F[1,14]=8.96, p=0.01).

One possible mechanism for tagging synapses for ingestion is via MFG-E8, a phosphatidylserine-recognising opsonin. Astrocytes, microglia and macrophages produce MFG-E8 both *in-vitro* and *in-vivo* which can bind to exposed phosphatidylserine on living neurons challenged with either Aβ or p-tau, facilitating phagocytosis via binding integrins on phagocytes ^29–33^. We have previously observed increased levels of MFG-E8 at the synapse in AD individuals compared to controls by un-biased proteomic screening ^34^. Here, we have validated that the AD synaptoneurosomes tested in the phagocytosis assays also express higher levels of MFG-E8 compared to control ones (p=0.01) by quantitative dot blots (Figure 6A-C). Thus, we tested the hypothesis that blocking the integrin binding site on MFG-E8 would modulate phagocytosis of AD and control synaptoneurosomes. Synaptoneurosomes from AD and control brain were pre-incubated with an antibody to block the integrin binding domain of MFG-E8 (or IgG1 isotype control) before applying them to cultured human microglia or astrocytes. In human astrocyte cultures (Figure 6D), anti-MFG-E8 antibody treatment significantly reduced phagocytosis of AD synaptoneurosomes (p=0.0011), compared to both untreated and IgG1 (isotype control for anti-MFG-E8 antibody) incubation (p=0.0384) (Figure 6E and Figure S5I). When astrocytes were exposed to control synapses, there was no effect of either MFG-E8 or IgG1 treatment on phagocytosis. This provides a novel mechanism by which astrocytes ingest AD-specific synapses and a pathway that can be targeted to modulate phagocytosis. In human brains, astrocytes abundantly express the integrin receptor α5β5 which is known to bind MFGE8 ^35^. Therefore, we used an antibody to block this receptor and test its effects in ingestion of synaptoneurosomes. We found that anti-α5β5 treatment on human astrocytes resulted in significantly reduced phagocytosis of AD (p=0.006), but not control (p=0.0758) synaptoneurosomes compared to untreated cells (Figure 6F). This suggests that the both MFG-E8 and its integrin receptor α5β5 are important molecular pathways in astrocytic ingestion of synapses in AD.

**Figure 6:**
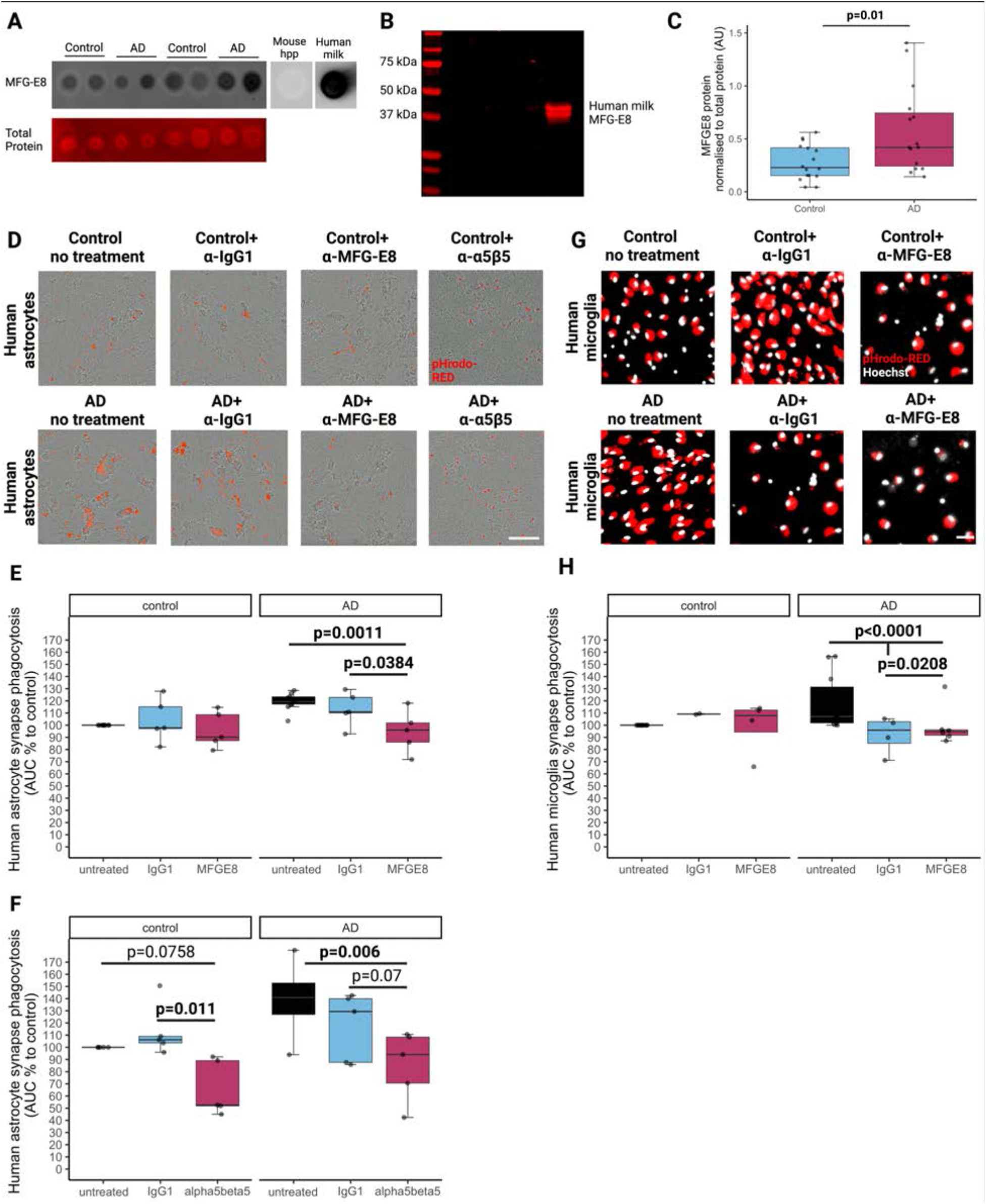
Synaptic MFG-E8 and astrocytic integrin α5β5 modulate phagocytosis of AD synaptoneurosomes. (A) Representative dot blots of aged control and AD-derived synaptoneurosomes probed with MFG-E8. Mouse hippocampus (hpp) was used as a negative control where no MFG-E8 binding is observed and human milk was used as positive control where MFG-E8 is highly expressed. (B) The anti-MFG-E8 antibody was validated by Western blot where it strongly binds to human milk at the correct predicted weight of 43kDa. (C) Quantification of dot blots in control (n=16) and Alzheimer’s disease (A)D) (n=15) synaptoneurosomes shows significantly higher levels of MFG-E8 in AD samples compared to control ones (ANOVA after linear mixed effects model on Tukey transformed data, F[1,29]=7.59, p=0.01). (D) Images from phagocytosis assays in human astrocytes treated with pHrodo-Red labelled human synaptoneurosomes from control and Alzheimer’s disease (AD) brains, illustrating differences in phagocytosis of synapses with pre-treatment of synaptoneurosomes with MFG-E8 antibody, integrin α5β5 antibody or IgG1 control antibody. Scale bar represents 200μm. (E) Area under the curve (AUC % normalised to last control for each experiment) of human astrocytes ingesting control or AD synaptoneurosomes shows statistically significant decrease in phagocytosis of treated with anti-MFG-E8 antibody, but not IgG1, compared to AD untreated synaptoneurosomes and a significant difference between IgG1 treated and MFG-E8 treated synapse phagocytosis (n=5 independent replicates; ANOVA after linear mixed effects model with disease status of synapse donor, time, and treatment as fixed effects and experimental replicate as a random effect shows an effect of disease of synapse donor F[1,30.2]=9.674, p=0.004058; treatment F[3,32.1]=237.4, p=2.2×10^-6^, and an interaction between disease status of synapse donor and treatment F[2,30.2]=4.63, p=0.0176. Post-hoc Tukey corrected tests show in the AD synapse treated cells that anti-MFG-E8 pre-treatment reduces phagocytosis compared to both no treatment (p=0.0011) and IgG1 treatment (p=0.0384). (F) Area under the curve (AUC % normalised to last control for each experiment) of human astrocytes ingesting control or AD synaptoneurosomes shows statistically significant decrease in phagocytosis of astrocytes treated with anti-α5β5 antibody, but not IgG1, compared to AD untreated astrocytes and a significant difference between IgG1 treated and anti-α5β5 antibody treated astrocytes (n=5 independent replicates; ANOVA after linear mixed effects model with disease status of synapse donor, time, and treatment as fixed effects and experimental replicate as a random effect shows an effect of disease of synapse donor F[1,20]=8.71, p=0.00788 and treatment F[1,20]=8.71, p=0.000234, but not a significant interaction between disease status of synapse donor and treatment F[2,20]=2.76, p=0.0872. Post-hoc Tukey corrected tests show in the AD synapse treated cells that anti-α5β5 pre-treatment reduces phagocytosis compared to no treatment (p=0.006) and non-significant reduction to IgG1 treatment (p=0.07). In control synaptoneurosome treated cells, there was a non-significant reduction in phagocytosis in anti-α5β5 pre-treatment (p=0.0758) but a significant reduction between IgG1 and anti-α5β5 pre-treatment (p=0.011). (G) Images from phagocytosis assays in human microglia treated with pHrodo-Red labelled human synaptoneurosomes from control and Alzheimer’s disease (AD) brains, illustrating differences in phagocytosis of synapses with pre-treatment of synaptoneurosomes with MFG-E8 antibody or IgG1 control antibody. Scale bar represents 20μm. (H) Area under the curve (AUC % normalised to last control for each experiment) of human microglia ingesting control or AD synaptoneurosomes show that none of the treatments significantly changes phagocytosis of synapses isolated from control brain. In contrast, both MFG-E8 and IgG1 treatment significantly rescued the amount of microglial phagocytosis back to control levels (ANOVA after linear mixed effects model with disease status of synapse donor, time, treatment, age of microglial donor, brain region donated, and sex of microglial donor as fixed effects and microglial donor ID as a random effect shows significant effects of treatment F[2,813.97]=11.69, p=9.84×10^-6^ and an interaction between disease status of synapse donor and treatment F[2,816.83]=39.19, p<2.2×10^-16^). Tukey post-hoc tests reveal significant differences in the AD group between no treatment and MFG-E8 antibody treatment (p<0.0001), no treatment and IgG1 treatment (p<0.0001) and between MFG-E8 and IgG1 treatment (p=0.0208). In the control synapse treated condition, there is a significant difference between untreated and IgG1 treated microglial phagocytosis (p=0.036) with IgG1 treatment causing a slight *increase* in phagocytosis of control synapses. Total n=11 donors with n=11 no treatment, n=6 MFG-E8 + no treatment, n=4 IgG1 + MFG-E8 + no treatment.

We also challenged human cultured microglia with synaptoneurosomes treated with the anti-MFG-E8 antibody or IgG1 isotype control (Figure 6G) as above. MFG-E8 antibody treatment reduced phagocytosis of AD synaptoneurosomes compared to untreated (p<0.0001) and IgG1 isotype control (p=0.0208). We also found that AD synaptoneurosome phagocytosis was decreased by IgG isotype alone (p<0.0001) compared to non-antibody-containing vehicle, despite prior blocking of low affinity Fc receptors (with anti-CD16/32) on microglia in all conditions (Figure 6H). Anti-MFG-E8 treatment did not affect microglial phagocytosis of control brain-derived synaptoneurosomes, similar to astrocytes (Figure 6H). The absence of an IgG1 effect in astrocyte cultures above is consistent with their lacking expression of Fc receptors. Overall, blocking synaptic MFG-E8 on AD synapses also reduced phagocytosis by human microglia.

## Discussion

In conclusion, we have demonstrated that microglia and astrocytes in human AD brains contain more synaptic protein than in control brains, and that *in-vitro* these cells preferentially ingest AD-derived synapses compared to controls. This builds upon previous evidence that microglia and astrocytes are capable of containing synaptic material in AD ^36–38^. We speculate that these differences in synaptic ingestion are due to AD-specific mechanisms, as in schizophrenia, a disease also characterised by reduced synaptic levels where microglia partake in increased synaptic phagocytosis^39^, we have found that there no differences in synaptic ingestion by microglia in post-mortem tissue of people with and without schizophrenia ^40^. In AD, this could in theory be beneficial in clearing dysfunctional or dead synapses or could be harmful in contributing to synaptic degeneration and removal of functional synapses. It is possible that both are true with a differing balance of beneficial and harmful contributions of glia to synaptic phenotypes at different stages of disease and in different parts of the brain. Indeed, there is evidence that when microglia phagocytose amyloid beta due to manipulation of TDP-43, synapses are also phagocytosed ^41^, which demonstrates that a potentially beneficial action of microglia, namely clearance of pathological proteins, may be linked to the potentially detrimental effect of synapse removal.

Here we have shown that MFG-E8 can contribute to the glial-mediated synapse loss observed in AD. MFG-E8 protein is likely increased on AD synapses due to binding phosphatidylserine, which is a well-known microglial opsonin that has been observed to play a role in neuron phagocytosis by microglia in mouse cultures ^29^. Phosphatidylserine performs many signaling roles and is a major component of all neuronal plasma membranes. *MFGE8* is expressed by astrocytes and microglia in cortical grey matter ^35,42,43^, while *ITGAV* and *ITGBV* are also expressed in these cell types, which is important as MFG-E8 is known to bind glial integrin α5β5 encoded by these genes ^44^. Notably in human brain, *ITGB3* was expressed only at very low levels in astrocytes and microglia indicating that the MFG-E8 effect we observe in human astrocytes is likely driven by interaction with integrin α5β5 not α5β3 which it can also bind ^35^. Published single nucleus sequencing data from Alzheimer’s disease vs control brain confirm that *MFGE8* is expressed in astrocytes and is significantly upregulated in AD vs control astrocytes ^43^. This suggests that AD brain-derived synapses already partially tagged with MFG-E8 as we showed previously ^34^ could be further opsonised by astrocytes secreting MFG-E8 during the culture thus reinforcing this pathway as a mechanism for astrocyte-mediated synapse uptake. *ITGB5* is expressed in microglia and astrocytes in these data and in AD it is upregulated in astrocytes but not microglia. Interestingly, in this dataset also showed that *ITGAV* is expressed in both microglia and astrocytes and is downregulated in AD in both cell types ^43^. *MFGE8*, *ITGB5*, and *ITGAV* mRNA are all found in nuclei as well as cell somata, making the data from single nuclear sequencing studies of human post-mortem brain likely a reliable reflection of the expression levels of these genes ^45^.

With a clearer understanding of glia-synapse interactions, there is potential to develop effective therapeutics to protect synapse function. Drugs that attenuate astrocytic and microglial phagocytic activity and anti-complement therapies have shown some potential to reduce AD-associated pathology in mice, notably limiting synapse loss and rescuing cognitive deficits ^13,19,46^. This is very important as it suggests glial cells do not only clear dystrophic or degenerating neurons ^37,38^, but may also eliminate healthy ones. Our data here show the relevance of this process in human AD brain and glial cells, and reveal mechanisms by which augmented AD synapse uptake can occur and that may be targeted for therapeutic manipulation.

## Acknowledgments

We wish to thank our human tissue donors and their families for making this work possible. We gratefully acknowledge Edinburgh Neuroscience and Dr. Jane Haley MBE for facilitating collaborations leading to this work. We also wish to thank the MRC Edinburgh Brain Bank for providing human tissue and the group of Prof. Nancy Ip, Prof. Oscar Harari, and Prof. Mark Fiers for sharing their single nucleus sequencing data freely with their publications. We would like to thank the Lothian Bioresource for facilitating biopsy tissue collection. We would like to thank Dr. Jason Early and Prof. David Lyons at CDBS and MS Society Edinburgh for assisting with Airyscan microscopy. Electron micrographs were captured with assistance from Dr. Jon Moss. We would like to thank Dr. Anisha Kubasik-Thayil at the IMPACT Imaging Facility for use of Imaris at the University of Edinburgh. Figures were created using Biorender.

## Funding

This work was funded by the UK Dementia Research Institute which receives its funding from DRI Ltd, funded by the UK Medical Research Council, Alzheimer’s Society, and Alzheimer’s Research UK, the European Research Council (ERC) under the European Union’s Horizon 2020 research and innovation programme (Grant agreement No. 681181), Alzheimer’s Research UK and the Scottish Government Chief Scientist Office (ARUK SPG2013-1), a Wellcome Trust-University of Edinburgh Institutional Strategic Support Fund, MND Scotland, and Alzheimer’s Society. Electron micrographs were obtained at the Roslin Institute 3D Electron Microscopy Facility (funded by BBSRC Institute Strategic Program support).

## Author contributions

Conceptualization: MT, MJDD, CMH, BWM, TLSJ

Methodology: MT, MJDD, CD, DK, PB, SM, BV, KP, AJS, JB, ED, JB, RKH, JT, CL, JK, DS, CS, VEM, RTK, GEH, CMH, PMB, BWM, TLSP

Investigation: MT, MJDD, CD, BV, KP, MH, ED, JB, MH, TLSP

Visualization: MT, TLSJ

Funding acquisition: MT, BWM, TLSJ

Project administration: BWM, TLSJ

Supervision: MT, MJDD, CMH, BWM, TLSJ

Writing – original draft: MT, TLSJ

Writing – review & editing: All authors

## Competing interests

TS-J is on the Scientific Advisory Board of Cognition Therapeutics and receives collaborative grant funding from 2 industry partners. None of these had any influence over the current paper. Remaining authors declare no competing interests.

## SUPPLEMENTARY FIGURES

**Figure S1.**
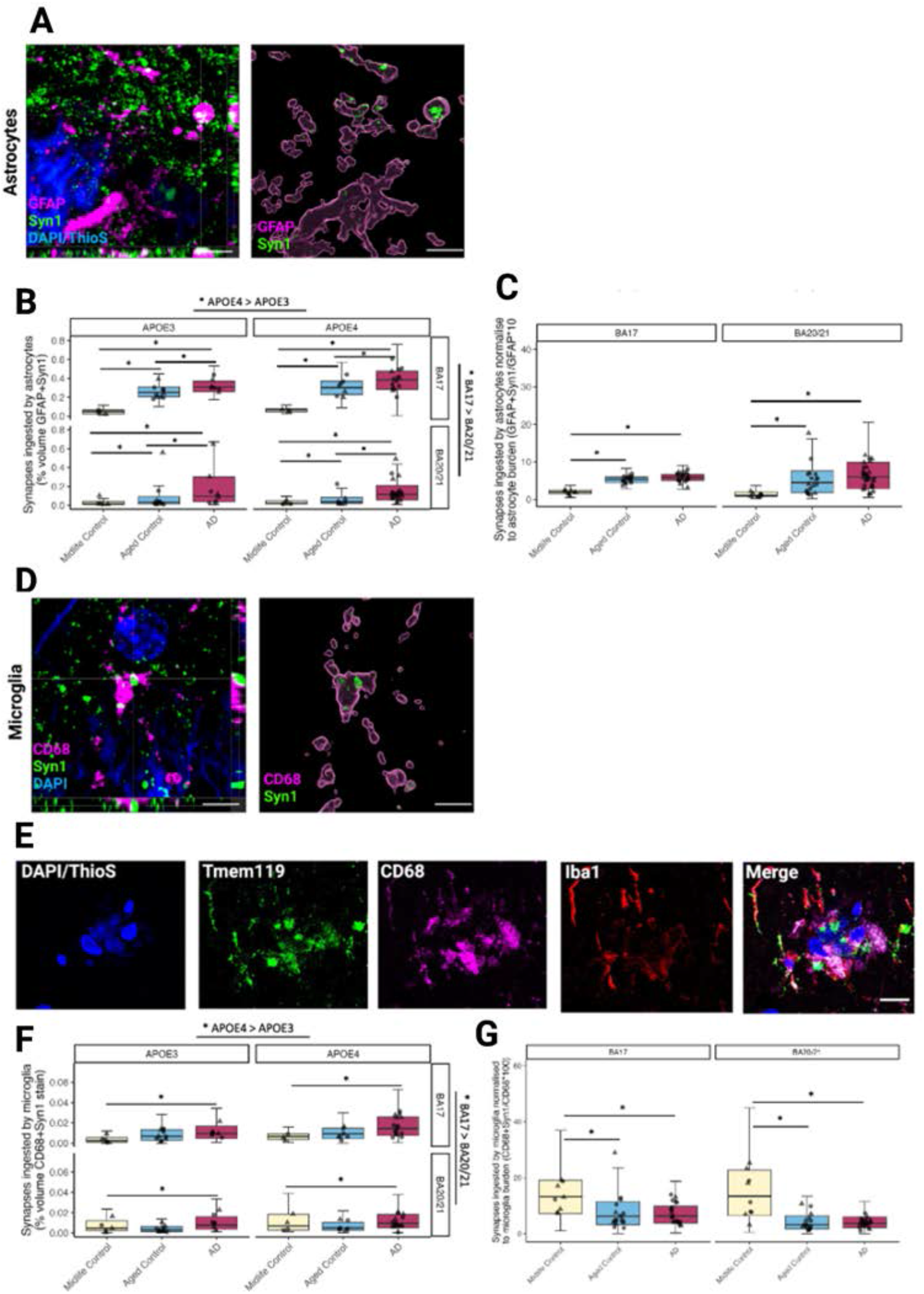
Supplementary statistics and validation microscopy. (A) Super-resolution confocal using an Airyscan microscope confirm synaptic engulfment by astrocytes in AD (orthogonal view left, 3D reconstruction right). Scale bar represents 5μm. (B) There is an effect of *APOE* genotype with *APOE4* carriers having more astrocyte engulfment than *APOE3* carriers. Also there is more engulfment in BA17 compared to BA20/21. (C) When normalized to GFAP burden, synaptic colocalization with GFAP was increased in AD compared to mid-life controls and increased in healthy ageing compared to mid-life controls. (D) Super-resolution confocal using an Airyscan microscope confirm synaptic engulfment by microglia in AD (orthogonal view left, 3D reconstruction right). Scale bar represents 5μm. (E) CD68 staining was confirmed to colocalise with Tmem119 and Iba1 (K). Scale bar represents 20μm. (F) There is an effect of *APOE* genotype with *APOE4* carriers having more astrocyte engulfment than *APOE3* carriers. Also there is more engulfment in BA17 compared to BA20/21. (G) When normalized to CD68 burden, the synaptic ingestion by microglia was no longer higher in AD cases compared to controls indicating the increase in AD in microglial ingestion of synapses is driven by microgliosis.

**Figure S2.**
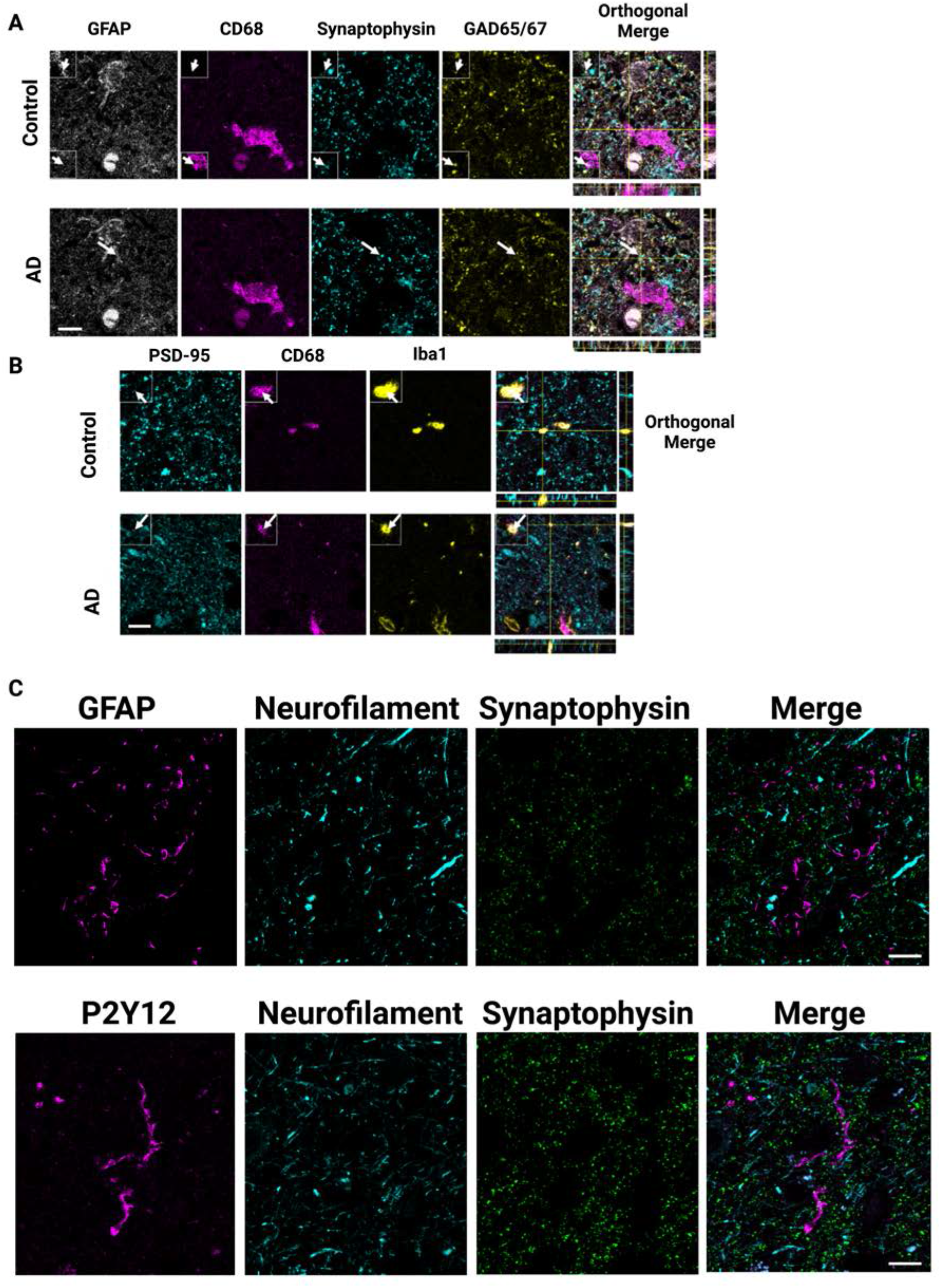
Glial co-staining with multiple synaptic markers and neurofilament. (A) Confocal images of staining with GFAP, CD68, synaptophysin, and GAD65/67 shows inhibitory synaptic protein inside astrocytes and microglia (arrows). (B) Excitatory postsynaptic protein PSD95 is also observed in CD68 and Iba1double positive microglia (arrows, B). Scale bars represent 10μm. Insets are 5 x 5 μm. (C) Astrocytes (GFAP) and microglia (P2Y12) do not overtly ingest axonal neurofilament in Alzheimer’s disease. Scale bars represent 10μm.

**Fig. S3.**
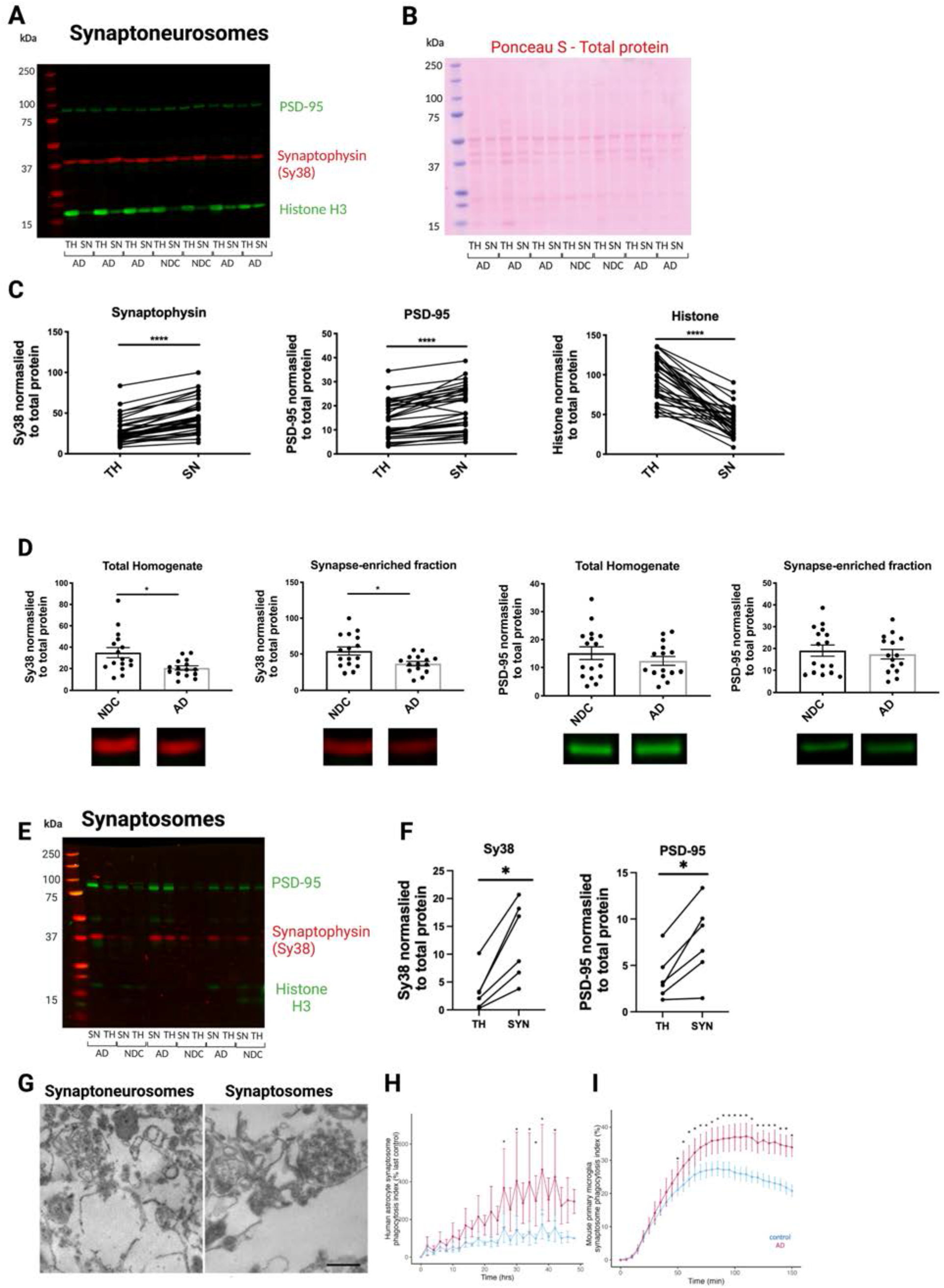
Validation of synaptoneurosome and synaptosome preparations. (A) Representative image of full-length Western blot, indicating whether a sample is from total homogenate (TH) or synaptoneurosomes (SN), and their corresponding disease status. Bands were quantified on Image Studio, and normalised to total protein, quantified by Ponceau S. (B) Ponceau S staining for total protein from gel shown in A (NDC=no disease control, AD=Alzheimer’s disease). (C) Significantly increased protein levels of the pre- and post-synaptic markers synaptophysin (Sy38) and PSD-95, respectively, as well as decreased protein levels of histone (H3), indicating exclusion of non-synaptic material (Wilcoxon matched-pairs signed rank test, ****p<0.0001, n=31). Lines link the two different preparations from the same case. (D) Decreased levels of synaptophysin (Sy38) in the total homogenate (Mann-Whitney test, p=0.0106) and synaptoneurosome fraction (unpaired Student’s t-test, p=0.0121) of AD cases (n=15), compared to age-matched NDC cases (n=16), as detected by Western blot. PSD-95 protein levels were not different between AD and NDC groups in total homogenate (unpaired Student’s t-test, p=0.332), nor in the synaptoneurosome fraction (unpaired Student’s t-test, p=0.627). (E) Representative image of full-length Western blot for synaptophysin, PSD-95 and histone H3, indicating whether a sample is from total homogenate (TH) or synaptoneurosomes (SN), and their corresponding disease status. Bands were quantified on Image Studio, and normalised to total protein, quantified by Ponceau S. (F) Significantly increased protein levels of the pre- and post-synaptic markers synaptophysin (Sy38) and PSD-95, respectively, (Wilcoxon matched-pairs signed rank test, *p<0.05, n=3). Lines link the two different preparations from the same case. (G) Synaptoneurosome and synaptosome pellets were embedded for electron microscopy confirming synaptic structures in both types of synaptic enrichment (scale bar 300 nm). (H) As seen with synaptoneurosomes, pHrodo tagged human synaptsomes from AD brain were phagocytosed more and faster by human astrocytes. (I) AD-derived human synapotosomes also ingested more than control ones by mouse microglia too, similar to the synaptoneurosomes. For statistics, *p<0.05.

**Fig. S4.**
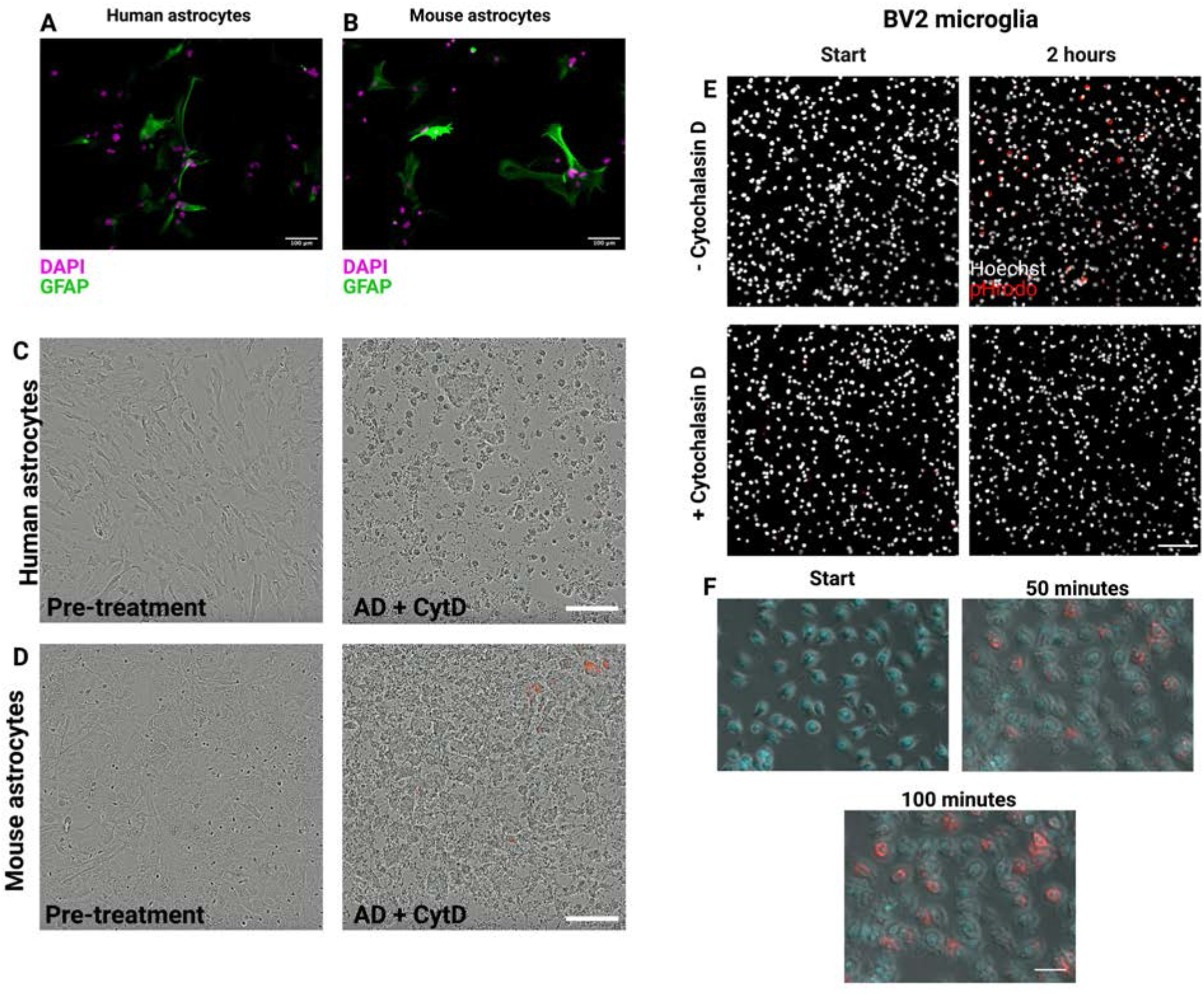
Validation of synaptoneurosome ingestion by astrocytes and microglia. (A) Primary human astrocytes stained for GFAP (green) shows most cells express the marker, indicating a pure astrocytic culture. (B) Primary mouse astrocytes stained for GFAP (green) shows cells express the marker, although some cells do not. Scale bars represent 100μm. (C) Primary human astrocytes treated with cytochalasin D prior to phagocytosis assay, blocking phagocytosis. Scale bar 200μm.(D) Primary mouse astrocytes treated with cytochalasin D prior to phagocytosis assay, blocking phagocytosis. Scale bar 200μm. (E) Still images from live imaging of BV2 microglia (Hoechst-positive nuclei in grey) undergoing phagocytosis of human synaptoneurosomes tagged with pHrodo. Synaptoneurosomes can be seen in red as they enter the acidic phago-lysosomal compartment of the cell. (F) Cells treated with 10μM of Cytochalasin D 30 minutes prior to the experiment showed no phagocytosis. Scale bar represents 30μm. (G) Live imaging of BV2 microglia (phase with Hoechst-positive nuclei in cyan) undergoing phagocytosis of human synaptoneurosomes tagged with pHrodo (can be seen as small spheroids on phase contrast). Synaptoneurosomes become red once they enter the acidic phago-lysosomal compartment of the cell. Each panel represents an image 50 minutes apart. Scale bar represents 50μm.

**Fig. S5.**
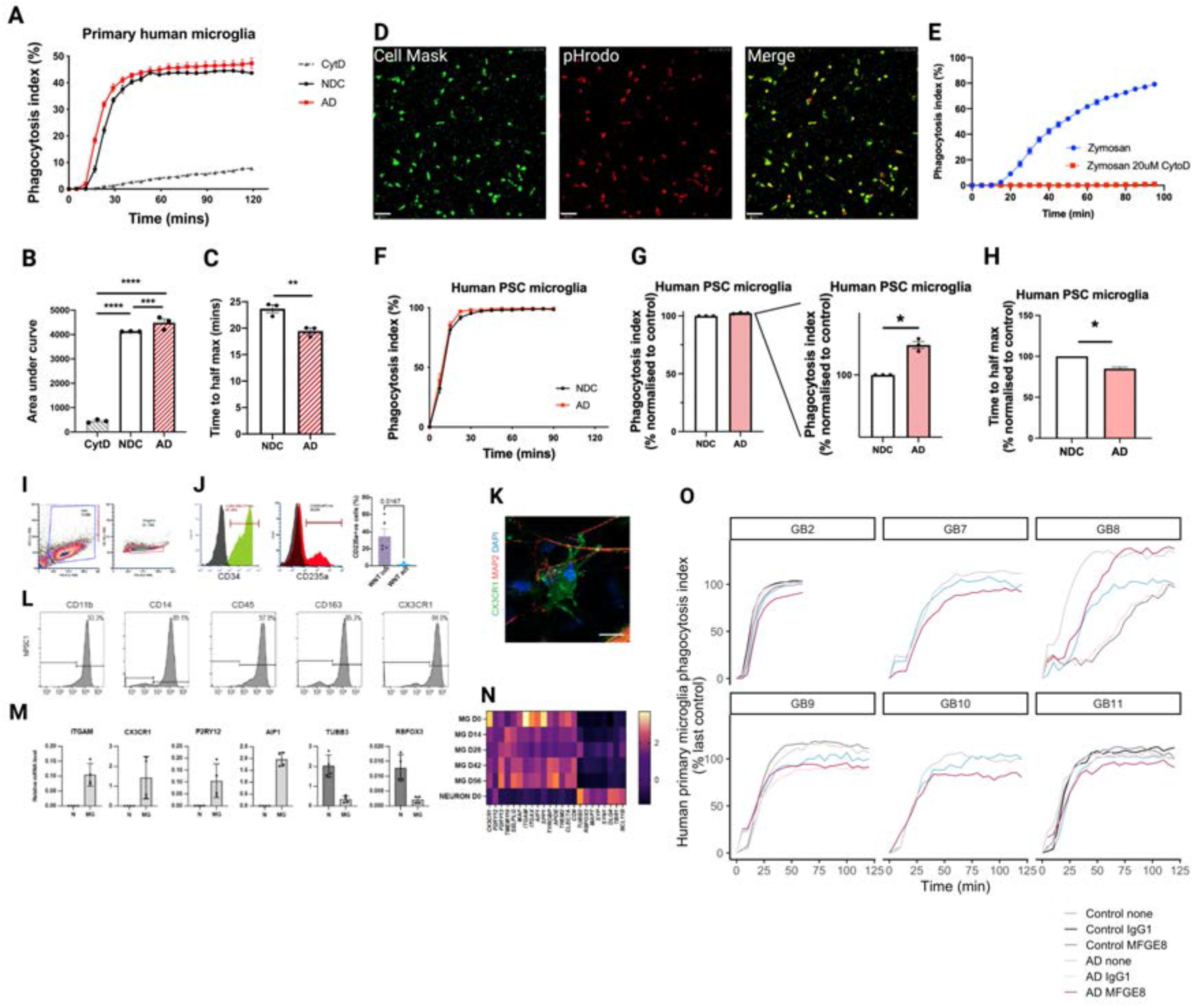
Increased phagocytosis of AD-derived synaptoneurosomes by primary human microglia (epilepsy). (A) Phagocytosis index of primary human microglia engulfing human synaptoneurosomes (n=1 human sample, replicated in 3 wells from an average of 9 images per well). CytD= cytochalasin D, NDC= non-demented control, AD= Alzheimer’s disease. (B) Area under curve from A (one-way ANOVA with Tukey’s multiple comparisons test, p=0.0004). (C) Time to half-maximum phagocytosis was calculated from the curve shown in A (unpaired Student’s t-test, p=0.0098). For statistics, **p<0.01, ***p<0.001, ****p<0.0001. Data shown as mean ± SEM. (D) Representative images of human PSC derived microglia-like cells labelled with Cellmask Deep red (green), showing phagocytosed pHrodo labelled synaptoneurosomes (red) at the end of imaging time. Scalebar: 80μm. (E) Zymosan beads are readily phagocytosed by human PSC microglia in culture, and treatment of cytochalasin D (CytoD) is sufficient to completely abolish it. Data shown as mean ± SEM. (F) All cell lines showed phagocytosis of SNS particles, the responsiveness and dynamics of microglia to AD and control material uptake depends on the individual cell lines. (G) Total amount of labelled particles as measured by the area under the curve was increased in PSC derived microglia presented with synaptoneurosomes from AD brain compared to NDC (one sample t-test, p=0.0142, hypothetical value=100). (H) PSC derived microglia-like cells phagocytosed AD SNS particles faster compared to NDC particles measured by the half max time (one sample t-test, p=0.016, hypothetical value=100). In all conditions, n=3 indicating lines from separate iPSC donors. For statistics, *p<0.05, and data shown as mean ± SEM). CytD= cytochalasin D, NDC= no disease control, AD= Alzheimer’s disease. (I) Human yolk sac haematopoetic precursors were differentiated from hIPSCs in 2 weeks. Floating single cells were analysed for markers of the lineage. (J) Yolk sac CD34+ precursor cells express CD235a marker transiently before differentiating into macrophages. Inhibition of WNT activity (purple) is necessary for yolk sac specific programme. (K) Yolk sac macrophages differentiate into CX3CR1+ microglia in MAP2+ neuronal co-cultures. Scale bar, 10μm. (L) Human iPSC derived microglia express lineage markers analysed by flow cytometry. (M) CD11B positive isolated microglia express high levels of mRNAs of lineage markers compared to neurons. Values are relative to YWHAZ. (N) Temporal pattern of microglia lineage markers *in vitro* reflects developmental changes *in vivo*. Values are Z-scores of mRNA levels. (O) The small surgical samples from which microglia were derived were not always sufficient to test all conditions. From our 11 donors, all had AD vs control synapse conditions, two had these and in addition AD synapses treated with MFG-E8 antibody, and 4 included all conditions (control and AD synapses with anti-MFG-E8, IgG1 control, or no treatment). In all 6 cases with MFG-E8 antibody pre-incubation, MFG-E8 antibody pre-treatment reduced AD synaptic phagocytosis compared to AD without antibody pre-treatment. In the 4 cases with IgG control, 2 had clear reduction in phagocytosis with MFGE8 compared to control IgG (GB11, GB2), one had no difference between MFGE8 and IgG (GB9), and one had increased phagocytosis with MFGE8 compared to IgG (GB8). From these data we conclude that while there is not a clear rescue of phenotype specifically with the MFG-E8 antibody treatment as seen for astrocytes, the microglial data show “responders” and “non-responders” to this treatment compared to IgG1 and all cases had a clear reduction in phagocytosis with either IgG1 or MFGE-8 treatment indicating that IgG on synapses may be an opsonin recognized by microglia.

**Figure S6.**
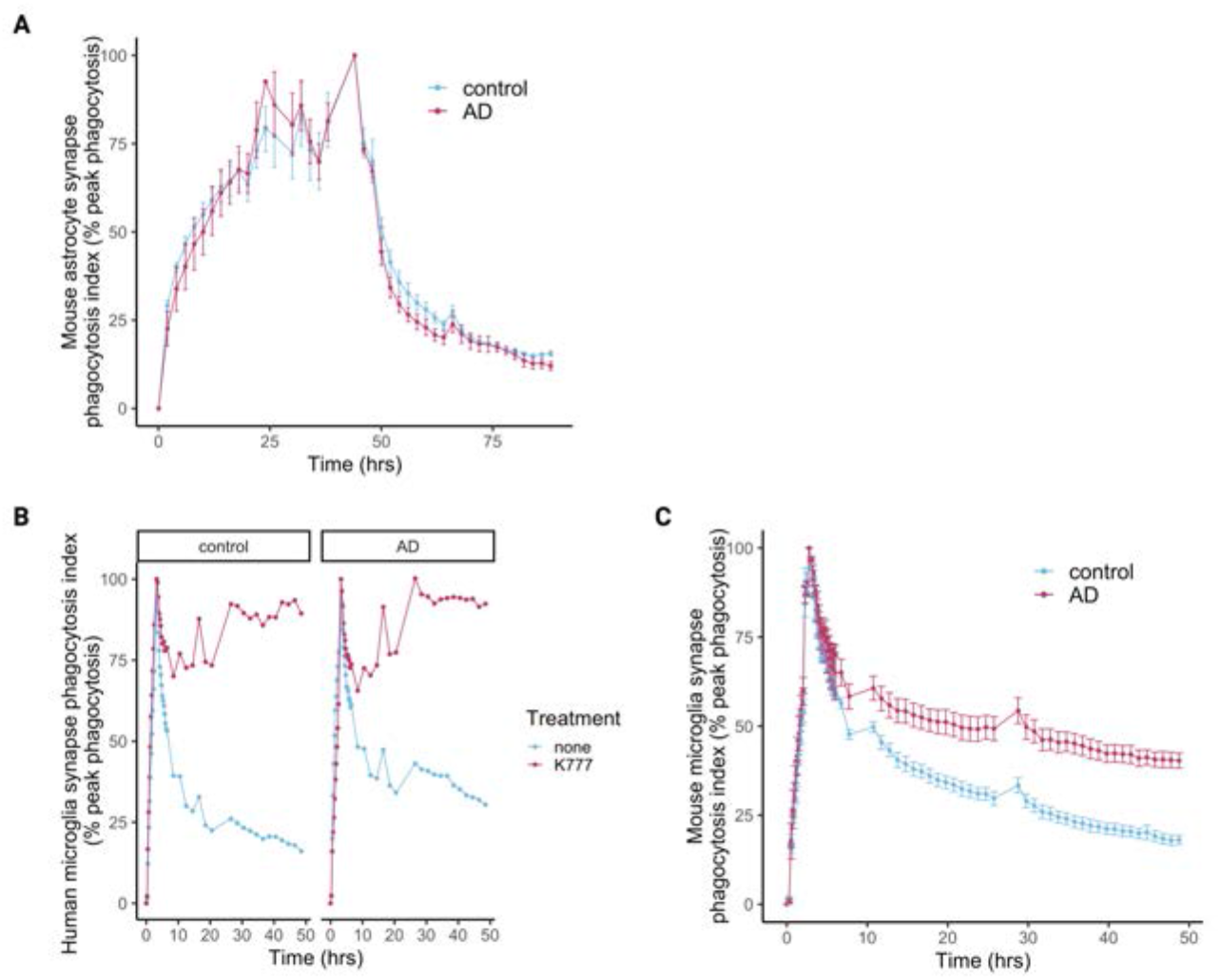
Degradation assays of mouse and human primary microglia. (A) Degradation assay in primary mouse astrocytes (n=4 replicates) of human control and Alzheimer’s disease (AD) synaptoneurosomes shows no differences in degradation throughout the assay. (B) Degradation assay in primary human microglia (n=1, GB12) of human control and Alzheimer’s disease (AD) synaptoneurosomes shows no differences in degradation in the 2 hours of the assay. Both control and AD synaptoneurosomes are efficiently degraded over time, which is blocked by the pan-cathepsin inhibitor K777. (C) Degradation assay in primary mouse microglia (n=4 adult mice) of human control and Alzheimer’s disease (AD) synaptoneurosomes shows no differences in degradation in the first 2 hours of the assay. Both control and AD synaptoneurosomes are efficiently degraded over time.

## STAR Methods

### RESOURCE AVAILABILITY

#### Lead contact

Further information and requests for resources and reagents should be directed to and will be fulfilled by the lead contact, Tara Spires-Jones.

#### Materials availability

This study did not generate new unique reagents.

#### Data and code availability

All spreadsheets of analysed data and statistical analysis scripts are available on the Edinburgh Data Share Repository (https://datashare.ed.ac.uk/handle/10283/3076) and as supplemental data files S1-S10. All image analysis scripts are available on Github at https://github.com/Spires-Jones-Lab. Raw images from analyses are available upon request from the lead contact.

### EXPERIMENTAL MODEL AND SUBJECT DETAILS

#### Animals

Microglial isolation experiments were performed using 12-week-old male C57Bl/6J mice (Charles River Laboratories). Mice maintained under a standard 12 h light/ dark cycle and provided with *ad libitum* access to food and water. Mice were housed in groups of up to five mice and were acclimatized for a minimum of 1 week prior to procedures. All experiments were conducted under the UK Home Office Animals (Scientific Procedures) Act 1986, in agreement with local ethical and veterinary approval (Biomedical Research Resources, University of Edinburgh).

#### Human post-mortem tissue

All tissue was provided by the MRC Edinburgh Brain Bank, following all appropriate ethical approval. For paraffin embedding, tissue was dehydrated via increasing ethanol solutions, fixed in formalin, and baked in paraffin-embedded blocks. Tissue from the inferior temporal lobe (Brodmann area 20/21) and primary visual cortex (Brodmann area 17) was cut using a microtome at 4μm thickness and mounted on glass slides for use in immunohistochemistry. The locations of these areas in the brain are visualised in Figure 1. AD cases were cross-checked neuropathologically and were confirmed to be Braak Stages V-VI. In one case (BBN: 24527) no plaques were measured in BA17 and was excluded as a whole from the near plaque analysis. Case BBN31495 has a Braak Stage of VI but has been cognitively tested upon 3 waves and was cognitively unimpaired. Data about subjects included in the study are found in Table S1. Use of human tissue for post-mortem studies has been reviewed and approved by the Edinburgh Brain Bank ethics committee and the ACCORD medical research ethics committee, AMREC (approval number 15-HV-016; ACCORD is the Academic and Clinical Central Office for Research and Development, a joint office of the University of Edinburgh and NHS Lothian). The Edinburgh Brain Bank is a Medical Research Council funded facility with research ethics committee (REC) approval (11/ES/0022).

#### BV2 microglia phagocytosis assay

Phagocytosis assays were optimised using the BV2 immortalised murine microglia cell line. BV2 microglia were cultured in DMEM + GlutaMAX (ThermoFisher Scientific, 31966-021) and were supplemented with 10% fetal bovine serum (FBS, ThermoFisher Scientific) and 1% penicillin/streptomycin (PenStrep, ThermoFisher Scientific). Cells were grown in a humidity-controlled incubator at 37°C with 5% CO_2_. One day prior to the phagocytosis assay, BV2 microglia were seeded in a 96-well flat bottom plate (ThermoFisher Scientific, 165305) at a density of 12,500 cells/well. Cells were stained with Hoechst (2μg/ml, ThermoFisher, H3570) to visualise nuclei and cytochalasin D treated cells (10μM, Sigma-Aldrich, C8273) were used as negative controls. BV2 cells were challenged with pHrodo-tagged synaptoneurosomes, and immediately taken to an ImageExpress high-throughput microscope (Molecular Devices) for live-imaging using a 20x air objective (37°C, 5% CO_2_). Images were taken every 5-10 minutes for 3 hours, using the same settings for all imaging sessions. All conditions were repeated in triplicate with 4 fields of view per well. For analysis, MetaXpress 6 (Molecular Devices) software was used to automatically calculate the number of cells per field of view and detect fluorescence around cells by thresholding. A phagocytosis index was calculated by normalising the number of cells phagocytosing to total number of cells in an automated and unbiased way. The same exposure settings were used for each set of experiments. Videos taken with phase contrast were recorded on a Zeiss Observer Z1 microscope using 20x air objective (37°C, 5% CO_2_), with images taken every 5 minutes.

#### Primary mouse microglia

Primary adult mouse microglia were isolated and cultured as described previously^47^. Brains from 12-week-old male C57/Bl6 (Charles-River) mice were isolated by terminally anaesthetizing with 3% isoflurane (33.3% O_2_ and 66.6% N_2_O) and transcardial perfusion with ice-cold 0.9% NaCl. Brains were immediately placed into ice-cold HBSS (ThermoFisher Scientific) and minced using a 22A scalpel before centrifugation (300 x g, 2 min) and digestion using the MACS Neural Dissociation Kit (Miltenyi) according to manufacturer’s instructions. Briefly, brain tissue was incubated in enzyme P (50 μL/brain) diluted in buffer X (1900 μL/brain) for 15 min at 37°C under gentle rotation before addition of enzyme A (10 μL/brain) in buffer Y (20 μL/brain) and further incubation for 20 min at 37°C under gentle rotation. Following digest tissue was dissociated mechanically using a Dounce homogenizer (loose Pestle, 20 passes) on ice and centrifuged (400 x g, 5 min at 4°C). To remove myelin, tissue was resuspended in 35% isotonic Percoll (GE Healthcare, GE17-0891-01), overlaid with 2 mL HBSS, and centrifuged (800 x g, 40 min, 4°C). Following centrifugation, the supernatant and myelin layers were discarded and the pellet resuspended in MACS buffer (PBS, 0.5% low endotoxin BSA (Sigma-Aldrich), 2 mM EDTA, 90 μL/brain). Anti-CD11b microbeads (Miltneyi) were added (10 μL/brain) and the suspension incubated for 15 min at 4°C. before running through pre-rinsed (MACS buffer) LS columns attached to a magnet (Miltenyi). After washing with 12 mL MACS buffer, columns were removed from the magnet and cells retained (microglia) were flushed in 5 mL MACS buffer. Microglia were resuspended in DMEM/F-12 (ThermoFisher Scientific) supplemented with 1% PenStrep, 10% FBS, 500 pg/mL rhTGFβ-1 (Miltenyi), 10 pg/μL mCSF1 (R&D Systems). Microglia were counted using a haemocytometer and plated out at 40,000 cells/well onto a black-walled, optical bottom 96-well plate (ThermoFisher Scientific) coated with poly-L-lysine. Cells were cultured for 7 days with a half media change on day 3. Phagocytosis assay was performed and analysed as described below in methods details with 9 fields of view per well.

#### Primary human (GBM) microglia isolation

Human microglial isolations were performed as described previously ^48^. Use of human brain tumour and peri-tumoural tissue resected at surgery for research was approved by National Health Service Lothian under protocol number LREC 15/ES/0094 issued to P.M.B. For isolation of human microglia, the same protocol was used as the mice (CD11b beads) with a few adjustments detailed as follows. For all steps until the Percoll gradient, RPMI (ThermoFisher, 11875093) with 3% FBS + 2 mM EDTA was used instead of HBSS. Additionally, as the tissue was not pefused, an additional treatment with red blood cell lysis buffer (Biolegend, 420301) was carried out following the Percoll gradient step. Microglia were counted using a haemocytometer and plated out at 40,000 cells/well onto a black-walled, optical bottom 96-well plate (ThermoFisher Scientific) coated with poly-L-lysine. Cells were cultured in DMEM/F-12 (ThermoFisher Scientific) supplemented with 1% PenStrep, 10% FBS, 500 pg/mL rhTGFβ-1 (Miltenyi), 10 pg/μL mCSF1 (R&D Systems) for 7 days with a half media change on day 3. Phagocytosis assay was performed and analysed as described above with 9 fields of view per well. Exclusion criteria for a case/donor was poor viability of the cells at the end of the experiment.

#### Primary human (epilepsy case) microglia isolation

Use of human temporal lobe resections for research was approved by National Health Service Lothian under protocol number 2017/0125/SR/720 issued to V.E.M. The protocol for isolating primary human microglia was adapted from a previously established one^48^. Fresh brain tissue was donated from a 23-year-old male undergoing epilepsy surgery. Resected brain specimen came from healthy tissue of the temporal lobe. Briefly, blood was removed by multiple washes of PBS followed by treatment with 0.25% trypsin and 100ug/ml DNAse in PBS for 30 minutes at 37°C, with gentle rotation. Trypsin was then deactivated with 10% FCS. Samples were centrifuged at 1,200 RPM for 10 minutes (high brakes) at 4°C and the supernatant was discarded followed by addition of PBS and Percoll (GE Healthcare) with further ultra-centrifugation at 15,000 RPM for 30 minutes at 4°C (no brakes). The myelin layer was aspirated off and cell layer was transferred in a clean tube, leaving behind a layer of red blood cells. The transferred cells were topped-up with PBS and centrifuged again at 1,200 RPM for 10 minutes (high brakes) at 4°C and resuspended in warm media containing 5% FCS and 0.1% glucose. Subsequently, isolated mixed cells were cultured in T12.5 flasks for 3 days. Three days later, microglia were trypsinised and collected from the flasks, following centrifugation to yield the microglia cell pellet. After the isolation, microglia were counted using a haemocytometer and plated out at 40,000 cells/well onto a black-walled, optical bottom 96-well plate (ThermoFisher Scientific) coated with poly-L-lysine, and were allowed to rest for 5 days prior to phagocytosis assays. Phagocytosis assay was performed and analysed as described above with 9 fields of view per well.

#### Human pluripotent stem cell derived microglia cultures

Human pluripotent stem cells were routinely maintained in mTESR media in GELTREX coated dishes. Pluripotent stem cells were differentiated to primitive macrophages using a modified protocol from Takata et al. (2017) [54]. Briefly, on day 0, PSC clumps were seeded on GELTREX coated dishes in mTESR media at 100,000 cells per 10cm^2^. Stempro-34 media (ThermoFisher Scientific, 10639011) was supplemented with 5 µM ascorbic acid (Sigma Aldrich, A5960), Glutamax (ThermoFisher Scientific, 35050038), penicillin and streptomycin (ThermoFisher Scientific, 15140122), apo transferrin (Sigma Aldrich, T1147), monothioglycerol (MTG) (Sigma Aldrich, M6145-25ml) and mixed fresh with growth factor combinations for every timepoint of the protocol. Floating cells were collected from day 8 and reseeded in fresh media after centrifugation. On day 16, media was switched to serum-free defined media consisting of IMDM (ThermoFisher Scientific, 31980030) and F12 (3:1 ratio) (ThermoFisher Scientific, 11765054) supplemented with 0.5% N2 (W-MRC CSCI in house) and 1% B27 supplement (ThermoFisher Scientific, 17504044) and BSA fraction V (0.1% g/100ml), penicillin and streptomycin and 50 ng/ml M-CSF (Miltenyi biotech, 130-096-493). Floating primitive macrophages were collected between day 28 and 35 and analysed for CD11b (CD11b-A594, Biolegend, 301340), CD45 (CD45-APC, Biolegend, 304011), CX3CR1 (CX3CR1-PECy7, Biolegend, 341611) expression by flow cytometry. Primitive macrophages were seeded on isogenic human cortical neuron cultures at 1:5 macrophage to neuron ratio in N2B27 co-culture media supplemented with 50 ng/ml IL34 (Peprotech, 20034205), 10 ng/ml M-CSF and 10 ng/ml BDNF (Peprotech, 45002100). Half the media was replaced with fresh media every 2-3 days. Microglia cells were collected from co-cultures after at least 2 weeks of coculture by magnetic immunolabelled sorting for CD11b+ cells (CD11b microbeads, Miltenyi biotech, 130-093-634). Purified microglia were replated on PDL coated Ibidi 8u slides in neuron conditioned co-culture medium supplemented with fresh cytokines with 5×10^4^ cells per well (1cm2). Cells were maintained for less than a week before imaging. Cellmask deep red (Life technologies, C10046) was used as contrast membrane labelling for 30 minutes before imaging. Imaging was done on Zeiss 710 with Ibidi incubator unit taking images from 10 regions of interest per condition. Image-series were analysed on Volocity software to measure pHrodo red intensity changes in cellmask deep red positive cells. Cell numbers measured for each cell line per condition.

#### Primary mouse astrocyte isolation

Astrocyte isolation and culturing has been previously described^49^. Glia cultures were achieved by passaging a mixed neuron/glia culture with trypsin consequently removing neurons. All reagents were purchased from Merk unless otherwise stated. Briefly, E17.5 mouse CD1 embryos were decapitated in accordance with schedule 1 of UK home office guidelines for humane killing of animals. The cortices were removed in a dissociation medium (81.8mM Na_2_SO_4_, 30 mM K_2_SO_4_, 5.84 mM MgCl_2_, 0.252 mM CaCl2, 1 mM HEPES, 20 mM D-glucose, 1 mM kynurenic acid, 0.001% Phenol Red) and then enzymatically digested with papain at 36,000 USP units/ml for 40 minutes. The cortices were then washed twice with dissociation medium followed by twice with plating media (DMEM+10% FBS+1X antibiotic-antimycotic agent (all Thermofisher)). Cortices were homogenized using a 5ml pipette by sequential suction/expulsion and plated a density of 2 cortices/T75 flask in plating medium. After 6-7 days *in vitro*, the mixed neuron/glia culture was trypsinized, centrifuged at 150g for 5 minutes and the pellet re-suspended in plating medium. 1/3 of the total cell suspension was plated into a new T75 which reached maximum confluency 4-5 days later. This cell population now predominately contains astrocytes with a smaller proportion of microglia which can be detached by shaking. Plating of astrocytes was done similar to microglia, as described above.

#### Primary human astrocyte line

Human astrocytes were purchased by Caltag Medsystems (SC-1800) and cultured in poly-D-lysine coated 96-well plates with Astrocyte medium (Caltag Medsystems, SC-1801), as described previously^50^. In previous work, the characterisation was confirmed by comparing genome-wide transcriptomic data and comparing it to published data relating to acutely purified human astrocytes, neurons, microglia and oligodendrocytes^50,51^. The expression of genes in primary human astrocytes (co-cultured with mouse neurons and rat microglia) significantly correlated with the expression in acutely isolated human astrocytes (*p>0.0001). Furthermore, genes predominantly expressed in acutely sorted human astrocytes compared to neurons, microglia or oligodendrocytes, were also significantly enriched in primary human astrocytes when compared to the other cell-types (*p>0.002).

### METHOD DETAILS

#### Immunohistochemistry for paraffin embedded human tissue

Paraffin-embedded sections were provided by the Sudden Death MRC Edinburgh Brain Bank and. Tissue was resected from post-mortem brains and dehydrated with ethanol, prior to paraffin-embedding. Sections were cut using a microtome at 4μm thickness and provided upon a justified tissue request. Slides with embedded tissue were dewaxed in xylene for 6 minutes, followed by rehydration using descending ethanol-to-water solutions: 100% ethanol, 90% ethanol, 70% ethanol, 50% ethanol, 100% water, of 3 minutes each. For antigen retrieval, samples were pressure cooked for 3 minutes at the steam setting in citrate buffer, pH 6 (Vector labs, H3300). Specifically, citric acid concentrate was diluted from 100x stock in de-ionised water for use, and made fresh each time. Slides were allowed to cool down under running water, and then immersed in 70% ethanol for 5 minutes. Slides were incubated with an autofluorescence eliminator reagent (Merck, 2160) for 5 minutes and washed with 70% ethanol, followed by two 5-minute washes using PBS-0.3% Triton X-100 (Sigma-Aldrich, T8787-100ML), and one wash with 1x PBS (Thermo Fisher, 70011036). Noise from red blood cells was eliminated using the Vector® TrueVIEW ® Autofluorescence Quenching Kit (SP-8400-15). Using a wax pen (Vector labs, H4000), the tissue was outlined and incubated with blocking solution for 1 hour. Blocking solutions consisted of 0.3% Triton X-100 and either 10% normal donkey serum (Sigma-Aldrich, D96663) or 5% normal donkey serum and 5% normal goat serum (Sigma-Aldrich, S26-100ML). Primary antibodies were diluted appropriately (Key Resources Table) at a final volume of 500μl per slide, and were allowed to incubate overnight (14-16 hours) in the cold room, at 4-6°C, in a humid chamber using wet paper towel in the staining box. On the second day of staining, slides were washed once in PBS-0.3% Triton X-100, followed by two 5-minute washes in PBS. Secondary antibodies were then applied at a volume of 500μl per slide (Key Resources Table). All cross-adsorbed secondary antibodies were made-up to a final concentration of 1:500 (4μg/ml) in PBS, and applied for 1 hour at room temperature. Nuclei were counterstained with DAPI (1μg/ml) (D9542-10MG, Sigma-Aldrich). For Thioflavin S (Sigma-Aldrich, T1892), slides were dipped in 0.001% Thioflavin S, made in 50% ethanol, for 8 minutes and differentiated in 80% ethanol for 1 minute. In the end, one drop of Immumount (Thermo Fisher, 9990402) was added per slide to allow for coverslip adherence. Coverslips size was No 1.5, corresponding to 22×40mm (VWR, 631-0136). Coverslips were pressed gently to remove excess mounting media and remove bubbles, and were allowed to dry at room temperature for at least 1 hour. Slides were kept in the fridge for long term storage and were allowed to reach room temperature prior to imaging.

#### Confocal microscopy and Image analysis

Slides were imaged on a confocal microscope (Leica TCS8) with a 63x oil immersion objective. Laser and detector settings were kept constant between samples. During imaging and analysis, the experimenter was blinded to brain area and disease status. Twenty images from the grey matter were taken per case, sampling randomly through all six cortical layers. The synaptic stain was used to confidently separate grey matter versus white matter. The resolution of the confocal microscopy was 0.18μm x 0.18μm x 0.3μm (xyz). Image stacks were segmented in 3D in Matlab under the auto-local threshold function with custom made scripts. For the CD68 stain, the Matlab segmentation setting were: window size=70, Factor C=0.2, method=mean, no minimum size selected. For the synapsin I stain the settings were: window size=10, Factor C=1, method=mean, no minimum size selected. For the GFAP stain, the Matlab segmentation setting were: window size=70, Factor C=0.15, method=mean, no minimum size selected. The same segmentation parameters were used for all images to keep the final outputs similar and not influence the final burdens between cases. Once the segmentation was completed, the images were processed with a custom Python script in order to obtain their volumetric measurements (i.e. density of objects/mm^3^ of tissue). This provided density measurements for stained markers, like the total objects of CD68-positive staining per image slice in a Z-stack. This allowed further analysis on Python for calculating the area of colocalisation between CD68/GFAP and SynI objects in 3D. To allow colocalization, two objects need to overlap at least 25% of their whole structure. Images were processed again with FIJI in order to add all segmented slices in their respective stacks in order to calculate the volume occupied by all of the staining in the stack. Each stack was normalised to the volume of the entire stack (different stacks have different amounts of slices, so the final volume differs). All scripts can be found in the following link: https://github.com/lewiswilkins/Array-Tomography-Tool.

#### Airyscan confocal microscopy

Images were acquired using a Zeiss LSM880 with Airyscan (Carl Zeiss Ltd, Cambridge, UK) point scanning confocal, fitted to an Axio Examiner Z1 microscope stand (Carl Zeiss Ltd, Cambridge, UK), running Zen Black 2.3. A 63x 1.4 NA oil immersion objective was used (Carl Zeiss Ltd, Cambridge, UK), with the Airyscan detector set to SR-mode. Images were acquired with optimal pixel size and z-interval.

#### Synaptoneurosome preparation

Preparation of synaptoneurosomes was performed as described previously ^24^. Snap-frozen human tissue of 300-500mg from BA38 (temporal cortex) was homogenised using a Dounce homogeniser with 1ml of a protease inhibitor buffer, termed here Buffer A. Buffer A consists of 25mM HEPES, 120mM NaCl, 5mM KCl, 1mM MgCl_2_, 2mM CaCl_2_, protease inhibitors (Merck, 11836170001) and phosphatase inhibitors (Merck, 524629-1SET) made up in sterile water, and was prepared fresh each day. The Dounce homogeniser and Buffer A were kept ice-cold throughout the procedure to avoid further degradation. We allowed 10 passes for full homogenisation but minimise cellular disruption that would lead to an impure final fraction. Once homogenised, the homogenate was aspirated in a 1ml syringe and passed through an 80-micron filter (Merck, NY8002500) to remove debris and yielded the total homogenate (TH). The filter was pre-washed with 1ml of Buffer A to maximise yield. A sample of the TH was snap-frozen on dry ice for Western blot analysis, and the rest was split in half for Western blot analysis and the phagocytosis assays. A subsequent filtration took place using a 5-micron filter (Merck, SLSV025LS) followed by centrifugation at 1000xg for 7 minutes to yield the synaptoneurosome (SN) pellet. In this step, extra care was taken to slowly pass the homogenate through the filter in order to prevent the filter breaking. The filter was pre-washed with 1ml of Buffer A to maximise yield. The pellet was washed with Buffer A and pelleted down again to ensure purity. Pellets were snap frozen on dry ice and stored at -80°C for long-term storage.

#### Synaptosome preparation

Synaptosome preparations were made as described previously^25^. Frozen human brains from BA38 and BA41/42 were homogenised in ice-cold Dounce homogenisers (10 passes) in 1ml of 0.32M sucrose solution (0.32M sucrose, 1mM EDTA, 5mM Tris-HCl, protease inhibitors (Merck, 11836170001) and phosphatase inhibitors (Merck, 524629-1SET), pH 7.4). The homogenate was spun at 900g for 4 minutes at 4oC and the supernatant was collected. The supernatant was spun again at 20,000g for 20 minutes at 4oC and the pellet was kept as the synaptosome fraction. Synaptosome pellets were snap-frozen in dry ice. A total homogenate aliquot was kept for Western blot validation of synaptic enrichment.

#### Electron microscopy

Synaptosome and synaptoneurosome pellets were fixed in 4% Paraformaldehyde, 2.5% Glutaraldehyde, 0.2% Picric acid in 0.1M phosphate buffer, pH 7.4 for 2 hours at room temperature plus overnight at 4 °C, post-fixed in 1% Osmium tetroxide for 30 minutes, washed in PB, boiled distilled water, and 50% ethanol, incubated in 1% uranyl acetate in 70% ethanol, dehydrated through 15 min steps in a graded series of ethanol then propylene oxide, 50% propylene oxide/50% Durcupan resin, then 100% Durcupan resin overnight in a Leica EM TP processor. Samples were baked in Durcupan resin in agar capsules overnight at 60 °C, cut on an Ultracut microtome (Leica) with a Histo Jumbo diamond knife (Diatome) into 70nm sections which were mounted on nickel mesh grids. Electron micrographs were captured on a Zeiss Gemini 360 scanning electron microscope with an annular STEM detector.

#### Protein Extraction

For protein extraction, samples we diluted five-fold with Tris-HCl buffer pH 7.6 (100mM Tris-HCl, 4% SDS, and Protease inhibitor cocktail EDTA-free [ThermoFisher Scientific, 78447]), followed by centrifuging for 25 minutes at 13,3000 RPM at 4°C. Then, the supernatant was collected in fresh tubes and heated at 70°C for 10 minutes. Samples were kept at -80°C for long-term storage.

#### Micro BCA

Micro BCA™ Protein Assay kit (ThermoFisher Scientific, 23235) was used to quantify protein levels, following manufacturer’s instructions. Briefly, 1 μl of sample was added in 1 ml of working solution and heated at 60°C for 1 hour. Working solution was made fresh right before required, consisting of 50% Buffer A, 48% Buffer B and 2% Buffer C, all provided in kit. Albumin was provided in the kit for determining a standard curve of the following concentration: 2 μg/ml, 4 μg/ml, 6 μg/ml, 8 μg/ml, 10 μg/ml, and 20 μg/ml. Absorbance values were obtained using spectrophotometry at 562 nm with working solution used as the blank value to calibrate the machine.

#### Western Blot

After protein extraction, each sample was made-up to 20 μg of protein as calculated by the micro BCA™ (protein extract diluted in de-ionised water) and diluted in half with Laemmli buffer (2x stock) (S3401-10VL). In each well, 15μl of sample were loaded in 4-12% Bis-Tris gels (ThermoFisher Scientific, NP0323BOX). Each gel was run with 5 μl of molecular weight marker (Licor, 928-40000) in the first well. Gels were washed with de-ionised water and diluted NuPAGE buffer (ThermoFisher Scientific, NP0002) (20x stock) to remove bits of broken gel and residual acrylamide. Western blot chambers were filled with diluted NuPAGE buffer (600 ml per chamber), making sure each chamber compartment was locked securely and there was no leaking. Gels were run at 80V for 5 minutes, 100V for 1.5 hours and 120V for 30 minutes. Then, using a scraper tool, gels were cracked open from their plastic casing and soaked in 20% ethanol for 10 minutes prior to transferring using the iBlot 2 Dry Blotting System (IB21001). Pre-packed transfer stacks containing a PVDF membrane (ThermoFisher Scientific, IB24002) were assembled as per manufacturers recommendations, and samples were transferred for 8.5 minutes at 25 V. After transferring, the PVDF membranes were stained for 5 minutes with Ponceau S in 5% acetic acid (P7170-1L), and washed 3 times with 5% acetic acid to stain for total protein. Ponceau S stained membranes were scanned and analysed on FIJI to obtain a measurement of total protein per sample. Ponceau S was washed out with PBS and the PVDF membranes were blocked using Odyssey blocking buffer (Licor, 927-40000) for 30 minutes, following overnight incubation with primary antibodies made up in Odyssey block with 0.1% Tween-20, at 4°C with gentle shaking. The following primary antibodies were used: Synaptophysin (mouse, Abcam ab8049, 1:1000), PSD-95 (rabbit, Cell signalling D27E11, 1:1000), Histone (rabbit, Abcam ab1791, 1:1000), and MFG-E8 (sheep, R&D Systems AF2767, 1:500). The next day, membranes were washed 6 times with PBS-0.1% Tween-20, and incubated with the following LI-COR secondary antibodies for 30 minutes: IRDye 680RD Donkey anti-Mouse IgG, highly cross-adsorbed (LI-COR, 925-68072, 1:5000) and IRDye 800CW Donkey anti-Rabbit IgG, highly cross-adsorbed (LI-COR, 925-32213, 1:5000). All gels were imaged using the same exposure times and intensities using a LI-COR Scanner. The images were then uploaded on ImageStudio for analysis. For each band, the same size box was used to ensure all samples are measured equally. Each sample was normalised to its corresponding value of total protein.

#### Dot blots

Equal protein concentrations of protein-extracted synaptoneurosome samples were applied on nitrocellulose membrane using a vacuum dot blotter (CSL-D48, Cleaver Scientific). Membranes were air-dried and blocked with Odyssey blocking buffer (LI-COR, 927-40000) for 30 minutes, followed by incubation with an anti-MFG-E8 antibody (sheep, R&D Systems AF2767, 1:500) and rabbit anti-goat HRP-tagged secondary antibody (Abcam ab6741, 1:5000). Membranes were processed with SuperSignal™ West Dura Extended Duration Substrate (ThermoFisher Scientific, 34075) for 5 minutes and imaged on a LI-COR Scanner under chemiluminescence settings. Total protein was measured with Revert 700 (LI-COR, 926-11021). For each blot, the same size box was used to ensure all samples are measured equally. Each sample was normalised to its corresponding value of total protein. Mouse hippocampus was used as a negative control and human milk was used as a positive control.

#### Synaptoneurosome and synaptosome labelling with pHrodo Red-SE

First, pHrodo™ Red-SE (ThermoFisher Scientific, P36600) was diluted with DMSO according to manufacturer’s instructions to reach a concentration of 10 mM. Synaptoneurosome and synaptosome pellets were resuspended in 100 mM sodium carbonate buffer pH 9, adapted from a previously described protocol^26^. Synaptoneurosomes were tagged with pHrodo Red-SE based on protein concentration, roughly at 4 mg/ml, as calculated by the micro BCA, and incubated at room-temperature under gentle shaking for 1 hour, covered in foil. Samples were centrifuged at 13,000 RPM for 10 minutes to obtain the labelled synaptoneurosome pellet, followed by 3 rounds of washing with PBS and centrifugation to wash out any unbound dye. The pHrodo-labelled synaptoneurosome pellets were then resuspended in 5% DMSO-PBS and aliquoted for storage at -80°C. Aliquots of pooled AD and NDC samples were prepared fresh.

#### IncuCyte phagocytosis assay on astrocytes

Due to the long image-acquisition times acquired for astrocyte phagocytosis assays, these experiments were performed in an IncuCyte S3 live-imaging system that is housed in a tissue-cultured incubator. Phase images were taken prior to synaptoneurosome addition to normalise ingestion for the confluency of astrocytes per well. Astrocytes were imaged every 2 hours with the 20X objective for up to 48hours on phase contrast and red channel to pick up the pHrodo-Red signal. To determine the amount of phagocytosis, the amount of pHrodo-Red was normalised to the respective phase contrast of the well at time = 0.

#### MFG-E8 and integrin (α5β5) antibody blocking *in vitro*

A recombinant mouse anti-human MFG-E8 blocking antibody (Mc3) (low endotoxin) was purchased by Creative Biolabs (FAMAB-0225CQ-LowE), along with an isotype control mouse IgG1 (MOB-065CQ). Both antibodies were used at a concentration of 10μg/ml (as suggested by the supplier). Integrin α5β5 was blocked at a concentration of 2.5μg/ml (R&D Systems, MAB2528). Synaptoneurosomes were incubated with the anti-MFG-E8 and IgG1 antibodies and cells were incubated with the anti-α5β5 and anti-IgG1 antibodies for 4 hours at 37°C prior to challenging *in vitro*. Glial cells were blocked with Fc block (human: Biolegend, 422302) (mouse: Biolegend: 101302) at a dilution of 1:1000 for 30 minutes prior to synaptoneurosome challenging.

#### Degradation assay

Primary mouse and human microglia were incubated with 2.5μl of pHrodo-tagged synaptoneurosome preparations from control and AD brains and were imaged for 2 hours in IncuCyte S3 every 15 minutes. The media was then aspirated and cells were washed with warm fresh media twice, before returning them to IncuCyte to resume imaging for another 2 days, imaging every 2 hours. Astrocytes were assessed similar by were imaged every 2 hours for 48 hours prior to washing. K777 (Adipogen, AG-CR1-0158) was added at a concentration of 1μM to block cathepsins and prevent degradation as control.

#### Immunofluorescence of cultured cells

Plated cells were fixed for 10 minutes with 4% PFA. Subsequently cells were washed with PBS (pH 7.4) and incubated with 0.1% TritonX-100 for 30 minutes. After washing, blocking solution was applied for 1 hour (10% NDS, 3% Bovine serum albumin [A8806-5G], 0.1% TritonX-100, 0.1% Tween-20 in PBS). Primary antibodies were diluted in blocking solution and used in the same concentrations as in the human paraffin tissue, and left overnight at 4°C. Cells were washed 3 times with PBS and secondary antibodies were applied for 30 minutes with DAPI (1 μg/ml). Secondaries and DAPI were washed, and cells were topped with PBS and kept in the fridge. Images were taken with 10x and 20x objectives using a Zeiss Axio Observer Z1 microscope.

#### Quantification and statistical analysis

Statistical analyses were performed in R Studio ^52^. For human post-mortem studies, differences in demographics between groups were tested with analysis of variance for continuous variables (age, PMI|) and with Fisher’s tests for factor data (*APOE* genotype, sex). To investigate differences between groups, linear mixed effects models were used with case or culture replicate as a random effect to account for multiple measurements from each case in order to avoid pseudoreplication of data while maximising statistical power ^53^. Fixed effects in the models were disease status (AD or control), Brain region, APOE4 status, sex, age and plaque proximity as relevant. Human post-mortem glial colocalization with synapse data were normalised by dividing the volume of colocalised stains by the total volume of glial stain in each stack in the “normalized” data in supplemental figure 1. Supplemental data statistics done on Prism 9. Full statistical details including data spreadsheets, statistical models, and results can be found as supplemental data files.

**Table S1.** Human cases used in the study. (NA=not available)

| BBN/CaseID | SD number | Disease | APOE | Age (years) | Sex | PMI (hrs) | Braak Stage | Experiment |
| --- | --- | --- | --- | --- | --- | --- | --- | --- |
| 29693 | SD002/17 | Midlife control | APOE4 | 49 | F | 94 | NA | IHC |
| 24479 | SD005/15 | Midlife control | APOE4 | 46 | F | 76 | NA | IHC |
| 33613 | SD014/18 | Midlife control | APOE3 | 46 | F | 99 | NA | IHC |
| 28959 | SD022/16 | Midlife control | APOE3 | 39 | M | 86 | NA | IHC |
| 30169 | SD022/17 | Midlife control | APOE4 | 48 | M | 58 | NA | IHC |
| 28960 | SD026/16 | Midlife control | APOE3 | 37 | F | 126 | NA | IHC |
| 34244 | SD036/18 | Midlife control | APOE3 | 49 | F | 69 | NA | IHC |
| 24342 | SD053/14 | Midlife control | APOE3 | 33 | M | 47 | NA | IHC |
| 29906 | SD013/17 | Midlife control | APOE4 | 51 | M | 52 | NA | IHC |
| 001.28793 | SD017/16 | Aged Control | APOE3 | 79 | F | 72 | 2 | IHC |
| 14395 | SD014/13 | Aged Control | APOE3 | 74 | F | 41 | NA | IHC + synaptoneurosomes |
| 19686 | SD063/13 | Aged Control | APOE3 | 76 | F | 75 | 1 | IHC + synaptoneurosomes |
| 20122 | SD003/14 | Aged Control | APOE3 | 59 | M | 74 | NA | IHC + synaptoneurosomes |
| 22612 | SD022/14 | Aged Control | APOE3 | 61 | M | 70 | NA | IHC + synaptoneurosomes |
| 26495 | SD024/15 | Aged Control | APOE3 | 78 | M | 39 | 1 | IHC + synaptoneurosomes |
| 28402 | SD051/15 | Aged Control | APOE3 | 78 | M | 49 | 1 | IHC + synaptoneurosomes |
| 28406 | SD001/16 | Aged Control | APOE3 | 79 | M | 72 | 2 | IHC + synaptoneurosomes |
| 28797 | SD025/16 | Aged Control | APOE3 | 79 | M | 57 | NA | IHC + synaptoneurosomes |
| 29086 | SD034/16 | Aged Control | APOE3 | 79 | F | 68 | NA | IHC + synaptoneurosomes |
| 32577 | SD002/18 | Aged Control | APOE3 | 81 | M | 74 | 2 | IHC |
| 001.34131 | SD029/18 | Aged Control | APOE4 | 82 | M | 95 | 4 | IHC |
| 15809 | SD029/13 | Aged Control | APOE4 | 58 | M | 90 | NA | IHC + synaptoneurosomes |
| 16425 | SD032/13 | Aged Control | APOE4 | 61 | M | 99 | NA | IHC + synaptoneurosomes |
| 20593 | SD006/14 | Aged Control | APOE4 | 60 | M | 52 | NA | IHC + synaptoneurosomes |
| 22629 | SD035/14 | Aged Control | APOE4 | 59 | F | 53 | NA | IHC + synaptoneurosomes |
| 29082 | SD031/16 | Aged Control | APOE4 | 79 | F | 80 | 3 | IHC + synaptoneurosomes |
| 31495 | SD043/17 | Aged Control | APOE4 | 81 | M | 38 | 6 | IHC + synaptoneurosomes |
| 001.30142 | SD020/17 | AD | APOE3 | 88 | F | 112 | 2 | IHC |
| 001.29081 | SD030/16 | AD | APOE3 | 90 | F | 110 | 2 | IHC |
| 15258 | SD026/13 | AD | APOE3 | 65 | M | 80 | 6 | IHC + synaptoneurosomes |
| 19595 | SD062/13 | AD | APOE3 | 87 | M | 58 | 6 | IHC + synaptoneurosomes |
| 19994 | SD002/14 | AD | APOE3 | 87 | F | 89 | 6 | IHC + synaptoneurosomes |
| 24527 | SD056/14 | AD | APOE3 | 81 | M | 74 | 5 | IHC |
| 28410 | SD005/16 | AD | APOE3 | 62 | F | 109 | 6 | IHC + synaptoneurosomes |
| 28771 | SD010/16 | AD | APOE3 | 85 | M | 91 | 6 | IHC |
| 32929 | SD012/18 | AD | APOE3 | 87 | F | 83 | 4 | IHC |
| 001.26500 | SD039/15 | AD | APOE4 | 81 | M | 83 | 6 | IHC |
| 001.26732 | SD048/15 | AD | APOE4 | 76 | M | 66 | 6 | IHC + synaptoneurosomes |
| 001.28796 | SD021/16 | AD | APOE4 | 60 | F | 54 | 6 | IHC |
| 001.29135 | SD027/16 | AD | APOE4 | 90 | M | 73 | 6 | IHC |
| 001.30883 | SD034/17 | AD | APOE4 | 61 | F | 69 | 6 | IHC |
| 001.30973 | SD039/17 | AD | APOE4 | 89 | F | 96 | 6 | IHC |
| 001.31499 | SD044/17 | AD | APOE4 | 85 | M | 78 | 6 | IHC |
| 001.33636 | SD017/18 | AD | APOE4 | 93 | M | 43 | 2 | IHC |
| 001.33698 | SD022/18 | AD | APOE4 | 90 | F | 76 | NA | IHC |
| 10591 | SD003/13 | AD | APOE4 | 76 | M | 76 | 6 | IHC +<br>synaptoneurosomes |
| 15256 | SD034/13 | AD | APOE4 | 60 | M | 28 | 5 | IHC |
| 15810 | SD018/13 | AD | APOE4 | 73 | F | 96 | 6 | IHC |
| 15811 | SD021/13 | AD | APOE4 | 81 | F | 41 | 6 | IHC |
| 19690 | SD064/13 | AD | APOE4 | 57 | M | 58 | 6 | IHC +<br>synaptoneurosomes |
| 20995 | SD019/14 | AD | APOE4 | 60 | M | 86 | 6 | IHC +<br>synaptoneurosomes |
| 23394 | SD038/14 | AD | APOE4 | 88 | F | 59 | 5 | IHC +<br>synaptoneurosomes |
| 24322 | SD049/14 | AD | APOE4 | 80 | M | 101 | 6 | IHC +<br>synaptoneurosomes |
| 24526 | SD055/14 | AD | APOE4 | 79 | M | 65 | 6 | IHC |
| 24668 | SD058/14 | AD | APOE4 | 96 | F | 61 | 6 | IHC |
| 25739 | SD014/15 | AD | APOE4 | 85 | F | 45 | 6 | IHC +<br>synaptoneurosomes |
| 26718 | SD040/15 | AD | APOE4 | 78 | M | 74 | 6 | IHC +<br>synaptoneurosomes |
| 29521 | SD035/16 | AD | APOE4 | 95 | M | 96 | 6 | IHC +<br>synaptoneurosomes |
| 29695 | SD004/17 | AD | APOE4 | 86 | M | 72 | 6 | IHC +<br>synaptoneurosomes |

**Table S2.** Data from human donors from glioblastoma surgeries.

| **CaseID** | **Age** | **Sex** | **Brain Region resected** |
| --- | --- | --- | --- |
| GB1 | 55 | M | parietal lobe |
| GB2 | 50 | F | temporal lobe |
| GB3 | 50 | M | frontal lobe |
| GB5 | 50 | F | parietal lobe |
| GB6 | 60 | F | parietal lobe |
| GB7 | 60 | F | frontal lobe |
| GB8 | 60 | F | frontal lobe |
| GB9 | 50 | F | frontal lobe |
| GB10 | 60 | M | frontal lobe |
| GB11 | 70 | M | frontal lobe |
| GB12 | 71 | M | parietal-occipital lobe |

## Notes

### Summary of Updates

This version of the manuscript was updated after 2 rounds of peer review to include substantial data on astrocytes and a mechanism that reduces glial phagocytosis of synapses from AD brain.

